# Paramagnetic spin labeling of a bacterial DnaB helicase for solid-state NMR

**DOI:** 10.1101/2021.09.14.460235

**Authors:** Johannes Zehnder, Riccardo Cadalbert, Maxim Yulikov, Georg Künze, Thomas Wiegand

## Abstract

Labeling of biomolecules with a paramagnetic probe for nuclear magnetic resonance (NMR) spectroscopy enables determining long-range distance restraints, which are otherwise not accessible by classically used dipolar coupling-based NMR approaches. Distance restraints derived from paramagnetic relaxation enhancements (PREs) can facilitate the structure determination of large proteins and protein complexes. We herein present the site-directed labeling of the large oligomeric bacterial DnaB helicase from *Helicobacter pylori* with cysteine-reactive maleimide tags carrying either a nitroxide radical or a lanthanide ion. The success of the labeling reaction was followed by quantitative continuous-wave electron paramagnetic resonance (EPR) experiments performed on the nitroxide-labeled protein. PREs were extracted site-specifically from 2D and 3D solid-state NMR spectra. A good agreement with predicted PRE values, derived by computational modeling of nitroxide and Gd^3+^ tags in the low-resolution DnaB crystal structure, was found. Comparison of experimental PREs and model-predicted spin label-nucleus distances indicated that the size of the “blind sphere” around the paramagnetic center, in which NMR resonances are not detected, is slightly larger for Gd^3+^ (~14 Å) than for nitroxide (~11 Å) in ^13^C-detected 2D spectra of DnaB. We also present Gd^3+^-Gd^3+^ dipolar electron-electron resonance EPR experiments on DnaB supporting the conclusion that DnaB was present as a hexameric assembly.

## Introduction

Solid-state NMR spectroscopy represents a versatile tool for the structural characterization of proteins as well as for probing their conformational dynamics[1, 2]. Technical advances in solid-state NMR spectroscopy, such as the availability of high magnetic-field strengths[3, 4], high rotation frequencies in magic-angle spinning (MAS) experiments [5, 6] as well as the development of highly efficient multidimensional radiofrequency pulse sequences, have enabled the routine assignment of uniformly ^13^C-^15^N labeled proteins with molecular weights up to ~30 kDa[7], and determination of the structures of small to medium-sized proteins (<100-150 amino acids per symmetric monomer). Structure determination mostly relies on the collection of a large number of distance restraints extracted from dipolar coupling-based NMR experiments and backbone torsion-angle restraints derived from the chemical shifts, e.g. via TALOS predictions[8]. Since the dipolar-coupling constant between two nuclei (denoted here by *i* and *j*), *D_ij_*, is inversely proportional to the third power of the distance separating the two nuclei, the detected spin pairs have short distances. Collection of long-range restraints (with |*i*-*j*| >4), with small *D_ij_* values, requires recording spectra with long mixing times. While for large proteins signal overlap in the spectra recorded at long mixing times can become problematic, the resolution can be improved in these cases by going to higher dimensions or by increasing the external magnetic-field strength[3, 4].

Pseudo-contact shifts (PCSs) or paramagnetic relaxation enhancements (PREs), which are caused by the interaction of nuclear spins with metal ions or radicals bearing unpaired electrons[9, 10], are a promising alternative to dipolar coupling-based distance restraints. Due to the much larger gyromagnetic ratio of electrons, PCSs and PREs can be observed over long distances (~20-25 Å, for some lanthanides even up to 40 Å[11]) providing long-range structural restraints. By contrast, typical dipolar nuclear distance restraints do not exceed 8 Å.

The relaxation properties of nuclei in a protein containing a paramagnetic center are affected by PREs[9, 12–14]. This relaxation phenomenon is treated within the Solomon-Bloembergen relaxation theory[15, 16] and originates from the modulation of the dipolar part of the electron-nucleus hyperfine coupling[15]. The mathematical expressions for the contribution of the PRE to the longitudinal (Γ_1_) and transverse (Γ_2_) relaxation rate constants in the solid state, respectively, are given by equations (1) and (2):

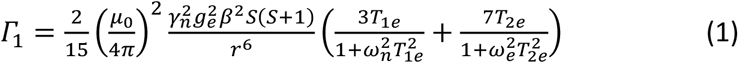

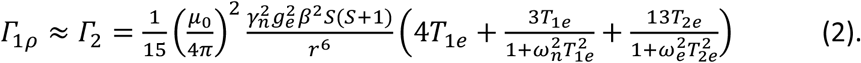

Herein, ω_n_ and ω_e_ represent the nuclear and electron Larmor frequencies. *T_1e_* denotes the longitudinal electron spin relaxation time constant with which the correlation time is approximated (with *T*_1e_ ≈ *T*_2e_)[17, 18] neglecting chemical-exchange effects.

Different sample preparation schemes for paramagnetic NMR studies have been described. In case of metalloproteins, naturally occurring metal binding sites can be used and loaded with transition metals or lanthanide ions[19]. For instance, the diamagnetic Mg^2+^ cofactor in ATP-driven proteins can be exchanged with paramagnetic metal ions[20, 21]. Additionally, diamagnetic proteins and nucleic acids[22] can be modified at specific positions with covalently bound paramagnetic tags[12, 23], e.g. by using tags that react with the thiol groups of free cysteines which can be site-specifically introduced into proteins by mutagenesis[24–27]. In case of membrane proteins, transition metal ions can be attached to the phospholipid head groups of the membrane by chelating tags[28–30]. Another alternative is to engineer completely new lanthanide binding sites into proteins[31-33].

PCSs and PREs can be integrated in both, resonance assignment and structure determination. The relative strengths of the two effects depend on the chosen paramagnetic tag or metal ion[10]. PREs and PCSs have been employed to support the 3D assignment of the uniformly or amino acid selectively labeled protein by comparing experimental PRE and PCS data with values calculated from the 3D structure, if the latter is known[34-40]. PCSs have been applied in various solid-state NMR studies[41-45] and PREs have been employed, for instance, in the structure determination of GB1[46-48].

We herein expand on these previous studies by applying paramagnetic tagging and solid-state NMR to the large bacterial DnaB helicase from *Helicobacter pylori*[49, 50]. DnaB is an ATP-fueled motor protein that unwinds double-stranded DNA into single strands during DNA replication[51]. In its active conformation, DnaB forms a ring-shaped hexamer with each polypeptide chain comprising 488 amino acid residues. The spin labeling of DnaB using a nitroxide (3-maleimido-PROXYL) and a lanthanide tag (Gd^3+^-maleimido-DOTA), respectively, is described. The efficiency of the labeling reaction was monitored by quantitative EPR spectroscopy carried out on the nitroxide-labeled protein. Site-specific PREs were extracted as signal attenuations from 2D ^13^C-^13^C Dipolar Assisted Rotational Resonance (DARR)[52, 53] and 3D NCACB correlation spectra using dia- and paramagnetic protein samples, for the latter without dilution with diamagnetic protein[42]. The PREs were compared to theoretical values. Those were obtained by generating ensemble distribution models of the nitroxide as well as the Gd^3+^ tag attached to the low-resolution X-ray structure of DnaB[54] by computational modeling with Rosetta[55]. Based on the structural models and their comparison with the experimental PRE data, we determined the blind sphere radii (i.e., the region in which NMR resonances are broadened beyond detection) for both tags. Finally, we present Gd^3+^-Gd^3+^ EPR distance measurements confirming that DnaB formed indeed a hexamer of dimers. Our study paves the way for combining paramagnetic solid-state NMR and EPR to unravel the interactions of DnaB with other proteins involved in DNA replication, such as the primase DnaG[56].

## Results and Discussion

### Spin labeling of the DnaB helicase with nitroxide and lanthanide tags

Spin labeling of DnaB was achieved by covalently modifying the two native cysteine residues in DnaB (C60 and C271) with either a nitroxide tag (3-maleimido-proxyl, abbreviated with PROXYL-M in the following) or a lanthanide-chelating tag (maleimido-DOTA, abbreviated with DOTA-M in the following) (Scheme 1, *vide infra*). The two tags were chosen in order to compare and contrast their applicability for protein solid-state NMR studies. The diamagnetic reference for the PROXYL-M-tagged protein was obtained by reducing PROXYL-M with ascorbic acid after tagging of DnaB[22]. In case of the DOTA-M tag, the paramagnetic protein sample was prepared by using Gd^3+^, whereas the diamagnetic reference state was obtained by using Lu^3+^.

**Scheme 1:**
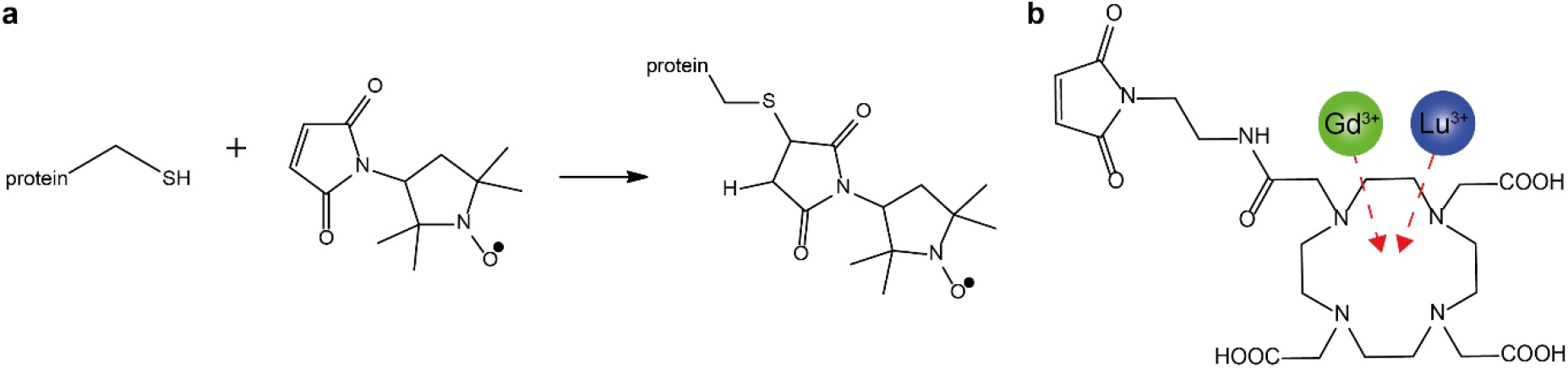
3-maleimido-proxyl (PROXYL-M) (**a**) and maleimido-DOTA (DOTA-M) (**b**) tags used for site-directed spin labeling of DnaB.

The crystal structure of *H. pylori* DnaB (resolution 6.7 Å) shows a stack-twisted double hexamer[54]. Each of the two hexamers can be described as a trimer of dimers which are assembled into a circular topology (see Figure 1a). The N-terminal domains (NTDs) and C-terminal ATPase domains (CTDs) of each DnaB monomer contribute to two separate rings, termed NTD- and CTD-rings, that have two different symmetries and give rise to a two-tiered hexamer (Figure 1b). While the NTD-ring is C3 symmetric and composed of a planar trimer of NTD dimers, the CTD-ring adopts a pseudo six-fold symmetry.

**Figure 1:**
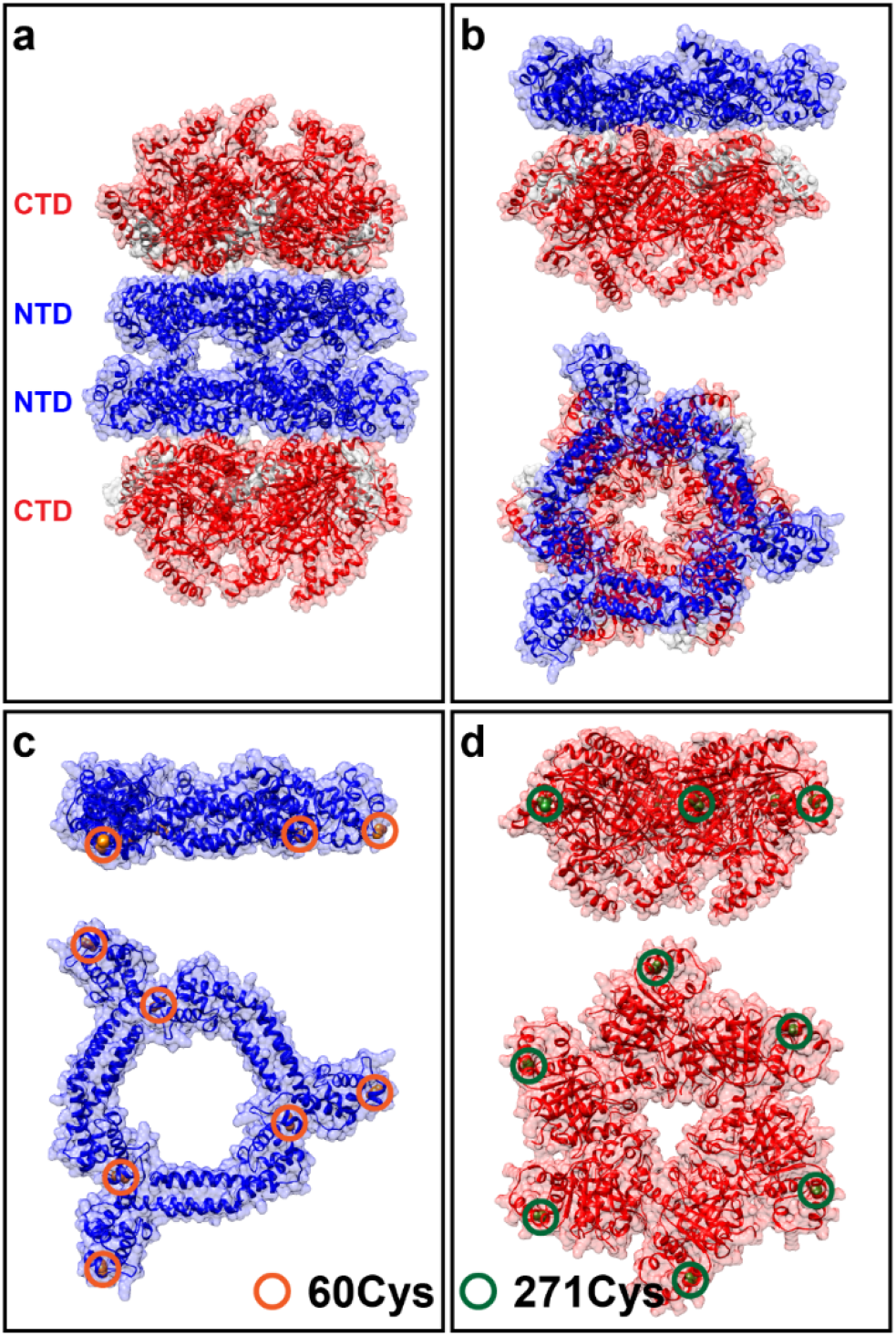
Spin labeling positions in DnaB. Dodecameric assembly of DnaB with the CTD colored in red and the NTD shown in blue (**a**), side and top view of one DnaB hexamer (**b**), side and top view of the NTDs within one hexamer with C60 highlighted by red circles (**c**) and side and top view of the CTDs within one hexamer with C271 highlighted by green circles. PDB ID 4ZC0 has been used to visualize the protein structure.

DnaB exhibits two cysteine residues per monomer which were employed for site-directed spin labeling. C60 is located in the loop following helix α3 in the NTD ring (Figure 1c), and C271 is located on helix α13 in the CTD ring (Figure 1d). Within the NTD homodimer, the two C60 residues experience different structural environments, one cysteine is oriented towards a neighboring DnaB subunit while the other cysteine is located on the outer surface of the NTD ring and is not involved in inter-subunit interactions. By contrast, within the CTD homodimer, the two copies of C271 have similar locations on the outer collar of the CTD-ring.

All four cysteine residues per DnaB homodimer are readily accessible to solvent. Within the DnaB crystal structure, the molecular surface area of the side-chain of these cysteine residues (determined using their atoms’ van der Waals radii) ranges from 87% to 95% relative to the surface area of the side-chain of the free amino acid. Thus, we expected nearly stoichiometric labeling efficiency of these sites when reacted with PROXYL-M and Gd^3+^-DOTA-M tags.

### Quantification of spin labeling efficiency by CW-EPR spectroscopy

NMR studies require labeling efficiencies of more than >80% to avoid any doubling of peaks owing to the presence of the resonances of both dia- and paramagnetic protein species in the NMR spectrum. This issue is of particular importance for large proteins because of the limited resolution of their NMR spectra. The success of the labeling reaction with the PROXYL-M tag was monitored by comparing the continuous wave (CW)-EPR spectra of unbound PROXYL-M dissolved in NMR buffer and of PROXYL-M-tagged DnaB solubilized in the same buffer (Figure 2a and b). As expected, the EPR resonance of PROXYL-M-tagged DnaB shows a significant anisotropic broadening pointing to slow molecule tumbling and thus an immobilized nitroxide species. Double integration of the EPR spectrum allowed determining the nitroxide spin concentration and calculation of the spin labeling efficiency of DnaB of 93±20% on average for the two labeling sites per DnaB monomer. Thus, we conclude that our spin labeling protocol is suitable to produce a nearly complete tagging of DnaB with PROXYL-M, which is required for solid-state NMR studies. Additionally, we assume from this result and from the accessibility of the cysteines that the labeling efficiency for the chemically similar DOTA-M tag will be in the same range as for PROXYL-M. The quantification of the Gd^3+^ spin concentration from CW EPR spectra is, however, challenging due to broad resonance lines owing to large zero field splittings.

**Figure 2:**
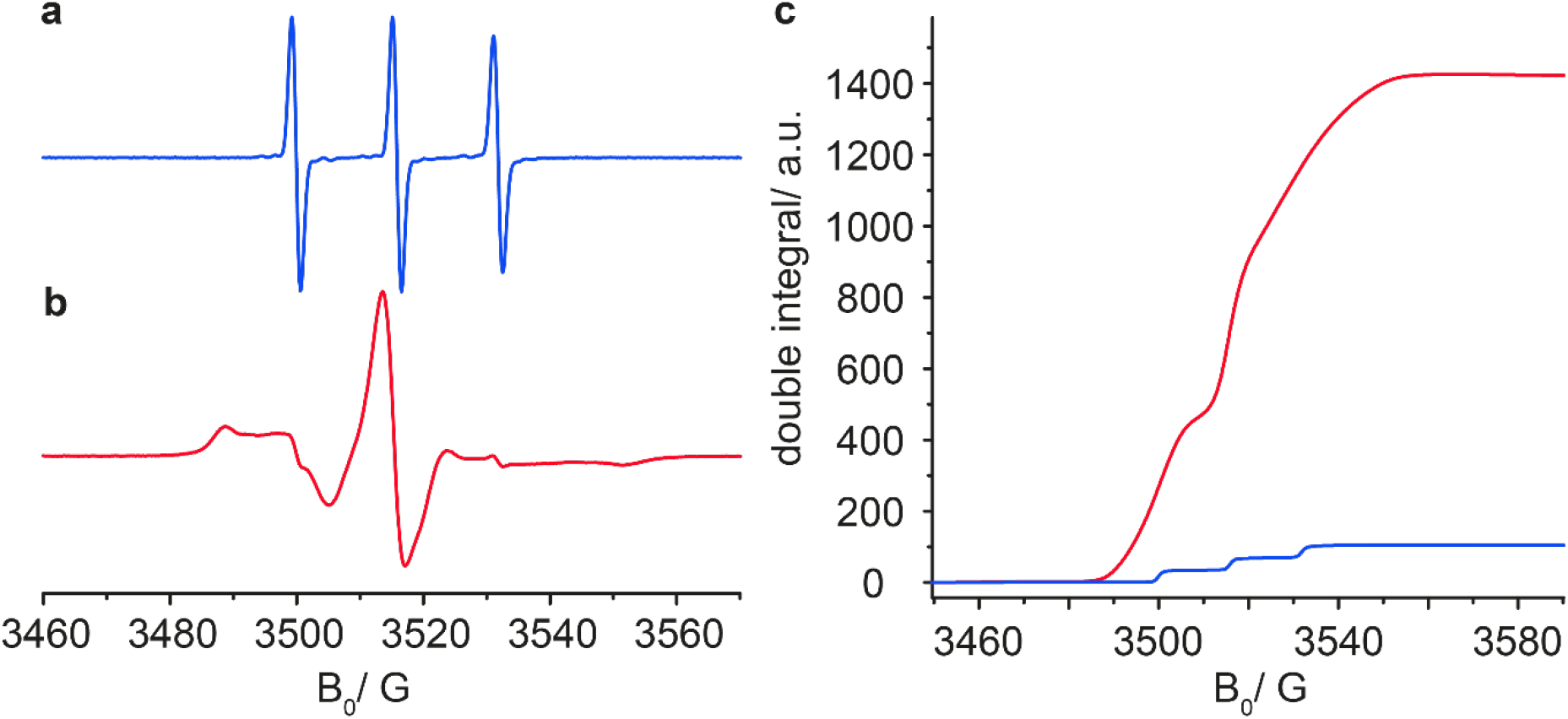
Determination of the spin labeling efficiency of PROXYL-M-tagged DnaB by EPR. CW-EPR spectrum of a 100 μM solution of free PROXYL-M (**a**, blue spectrum) and CW-EPR spectrum of PROXYL-M-tagged DnaB (738 μM protein concentration) (**b**, red spectrum). The double integration of both spectra (same color code as in **a** and **b**) is shown in **c**. Taking the concentrations into account, the spin labeling efficiency is determined to be 93 ± 20% on average for the two labeling sites per DnaB monomer.

### Solid-state NMR spectra of spin labeled DnaB

Figure 3 shows spectral fingerprints of the ^13^C-^13^C Dipolar Assisted Rotational Resonance (DARR) spectra recorded on DnaB tagged with PROXYL-M (red) and DnaB with the reduced tag (blue). Note that the paramagnetic sample was not diluted with diamagnetic protein. This is typically done for protein preparations in the microcrystalline state to suppress intermolecular paramagnetic effects[42]. However, we assumed that the impact of such intermolecular paramagnetic effects on the NMR spectrum is negligible for a sedimented protein sample, because in this case the protein molecules have no preferred orientation relative to each other. Structural details about sediments are still missing until today and more thorough investigations of such states are needed. Dilution of the paramagnetic protein with non-labeled diamagnetic protein has also practical limitations as this would further reduce the spectral signal-to-noise ratios precluding investigations of large proteins, such as DnaB.

**Figure 3:**
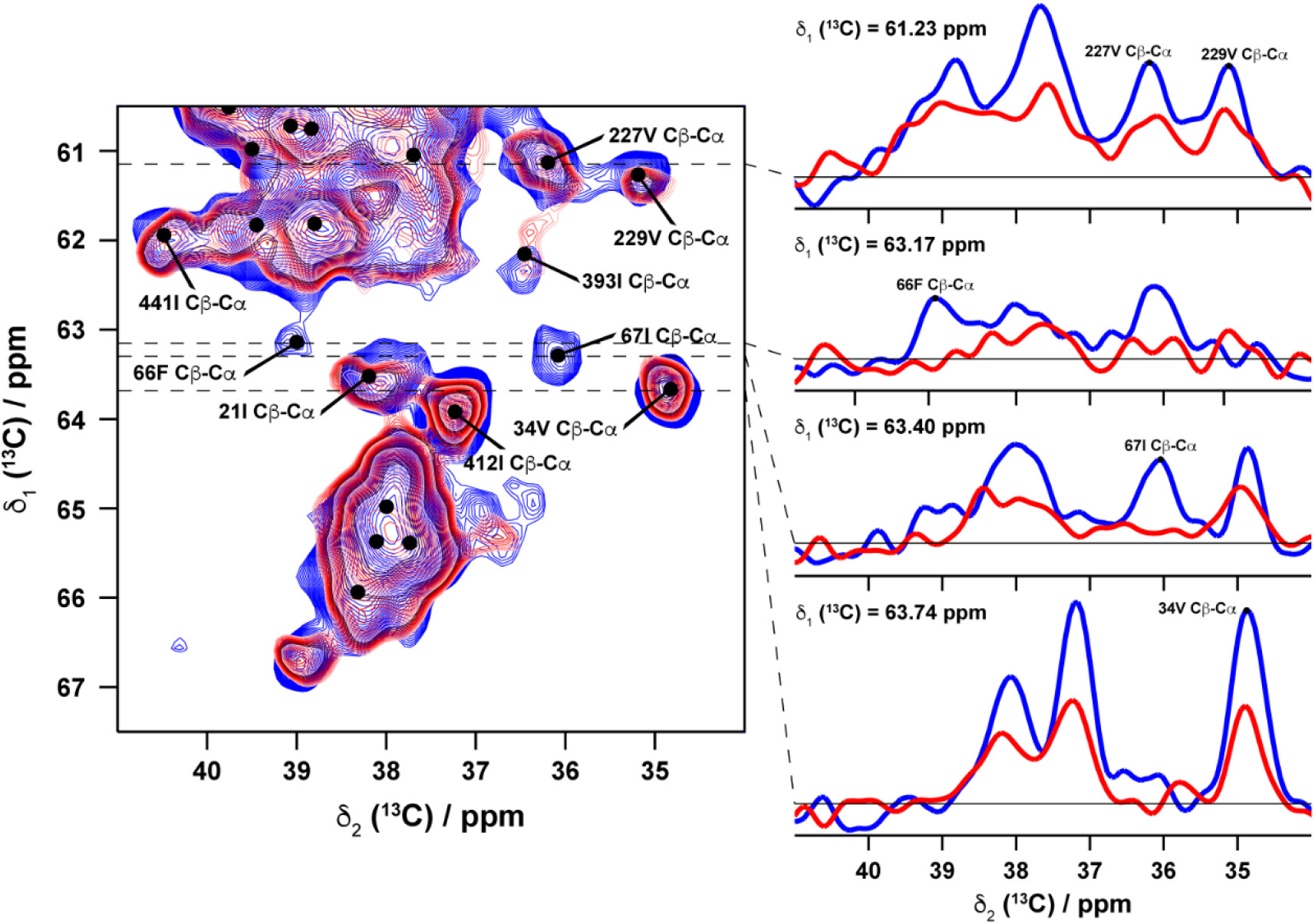
Detection of PREs in PROXYL-M-tagged DnaB by 2D DARR spectroscopy. Spectral fingerprints of ^13^C-^13^C DARR correlation spectra of PROXYL-M-tagged DnaB (red) and DnaB with reduced PROXYL-M tag (blue). 1D traces along the dashed lines are shown next to the 2D spectra. Complete DARR spectra of DnaB are shown in Figure S1 in the supporting material.

The absence of several resonances in the spectrum of PROXYL-M-tagged DnaB, e.g., of F66 and I67, is attributed to pronounced transverse PREs experienced by these residues. This can also be seen in the 1D traces along F2 (Figure 3) in which intensity reductions of additional peaks become apparent. Additional sections of the 2D DARR spectra showing the complete aliphatic region and another spectral fingerprint region, respectively, are displayed in Figures S1 and S2 in the supporting material. PREs were determined by comparing the intensities (peak heights) of the resonances in the spectrum of DnaB with the reduced PROXYL-M-tag (dia) with those in the spectrum of PROXYL-M-tagged DnaB (para); in the following expressed as the ratio I_para_/I_dia_ (note that the intensities were normalized to the correlation of D39).

Similarly, solid-state NMR spectra of DOTA-M-tagged DnaB were recorded. Spectral fingerprints are given in Figure 4. For the spectra showing the complete aliphatic region and another fingerprint region, respectively, see Figures S2 and S3. The spectrum of DOTA-M-tagged DnaB loaded with diamagnetic Lu^3+^ (reference sample) is shown in gray whereas the spectrum of DnaB tagged with paramagnetic Gd^3+^-DOTA-M is colored in orange. As for the PROXYL-M-tagged DnaB sample, pronounced PREs were observed in the spectrum of Gd^3+^-DOTA-M as cancellation of several NMR signals suggesting that labeling with Gd^3+^ was also nearly complete.

**Figure 4:**
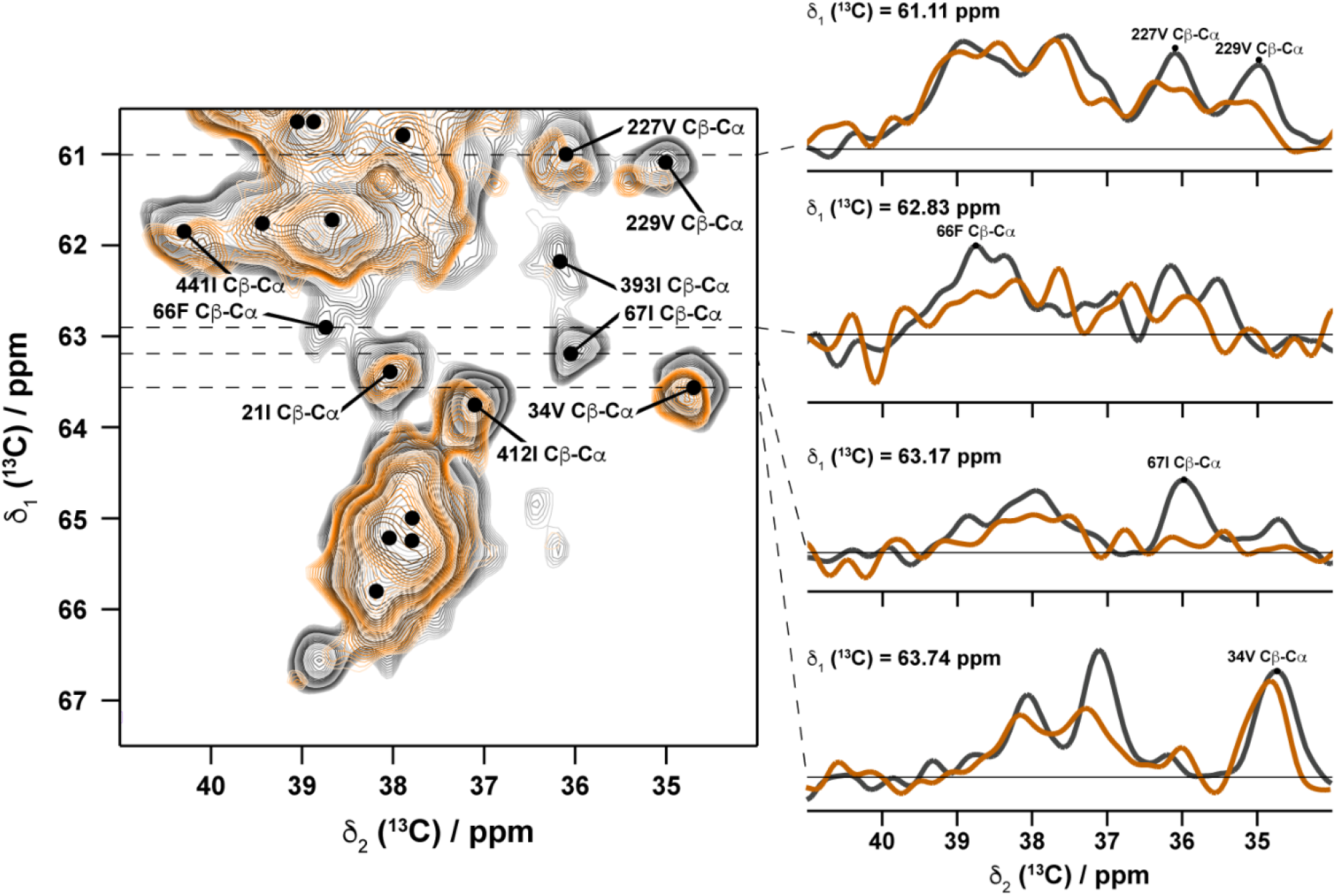
Detection of PREs in Gd^3+^-DOTA-M-tagged DnaB by 2D DARR spectroscopy. Spectral fingerprints of ^13^C-^13^C DARR correlation spectra of diamagnetic Lu^3+^-DOTA-M-tagged DnaB (gray) and paramagnetic Gd^3+^-DOTA-M-tagged DnaB (orange). 1D traces along the dashed lines are displayed next to the 2D spectra.

### Location of site-specific PREs in spin labeled DnaB and comparison with the DnaB crystal structure

For PROXYL-M-tagged DnaB, strong relaxation enhancements (I_para_/I_dia_ ≤ 0.25) of the NMR signals of D59, C60, P61, I62, F66, I67, T248, and C271 were observed. Residues D59-I62 are direct neighbors of C60 within the same polypeptide chain of DnaB. T248 on helix α11 is spatially close to C271 in the DnaB crystal structure model, which was initially derived from a low-resolution electron density (resolution 6.7 Å)[54]. Similarly, for Gd^3+^-DOTA-M-tagged DnaB, residues with pronounced PREs (I_para_/I_dia_ ≤ 0.25) included, besides C60 and C271, those residues which are in close proximity to the spin labeling sites in the DnaB crystal structure, namely D59, P61, I62, F66, I67, and T248.

In order to more quantitatively compare our experimental PRE data with the crystal structure model of DnaB, we used a computational modeling approach. We created ensemble conformation models[57] for PROXYL-M and Gd^3+^-DOTA-M and determined effective distances between the spin labels and every amino-acid residue in DnaB. First, we generated libraries of conformations for PROXYL-M and Gd^3+^-DOTA-M tags linked to the side-chain of Cys by sampling rotamers of small molecule fragments that matched regions in PROXYL-M-Cys or Gd^3+^-DOTA-M-Cys from the Crystallography Open Database[58, 59] (COD) (Figure S4). Based on the resulting conformer libraries, we then modeled the conformational ensembles of PROXYL-M and Gd^3+^-DOTA-M spin labels at every cysteine in the DnaB dodecamer using the Rosetta program[55] as basis for distance measurements.

Figure 5a shows the positions of the nitroxide O1 atom obtained from the distributed conformer models of PROXYL-M. Figure 5b displays the resulting O1 atom density at each of the four spin label sites within one DnaB homodimer. We determined effective spin label-amino acid residue distances by first calculating the O1 atom center of mass at every spin label site in the DnaB dodecamer and second measuring all pairwise distances between the centers of mass and the Cα or side-chain carbon atoms of residues in the asymmetric DnaB homodimer (see Materials and Methods). Figures 5c and 5d compare the experimentally observed PRE data for PROXYL-M-tagged DnaB with expected PREs that were calculated based on the effective spin label-amino acid distances. Overall, we found a favorable correlation between experimental and calculated PREs; for 82% of PREs the difference between experimental and predicted value was smaller or equal ±0.2. This supports the notion that our computational models of the ensemble of PROXYL-M spin labels based on the low-resolution DnaB crystal structure provide a faithful atomistic interpretation of the experimental NMR measurements.

**Figure 5:**
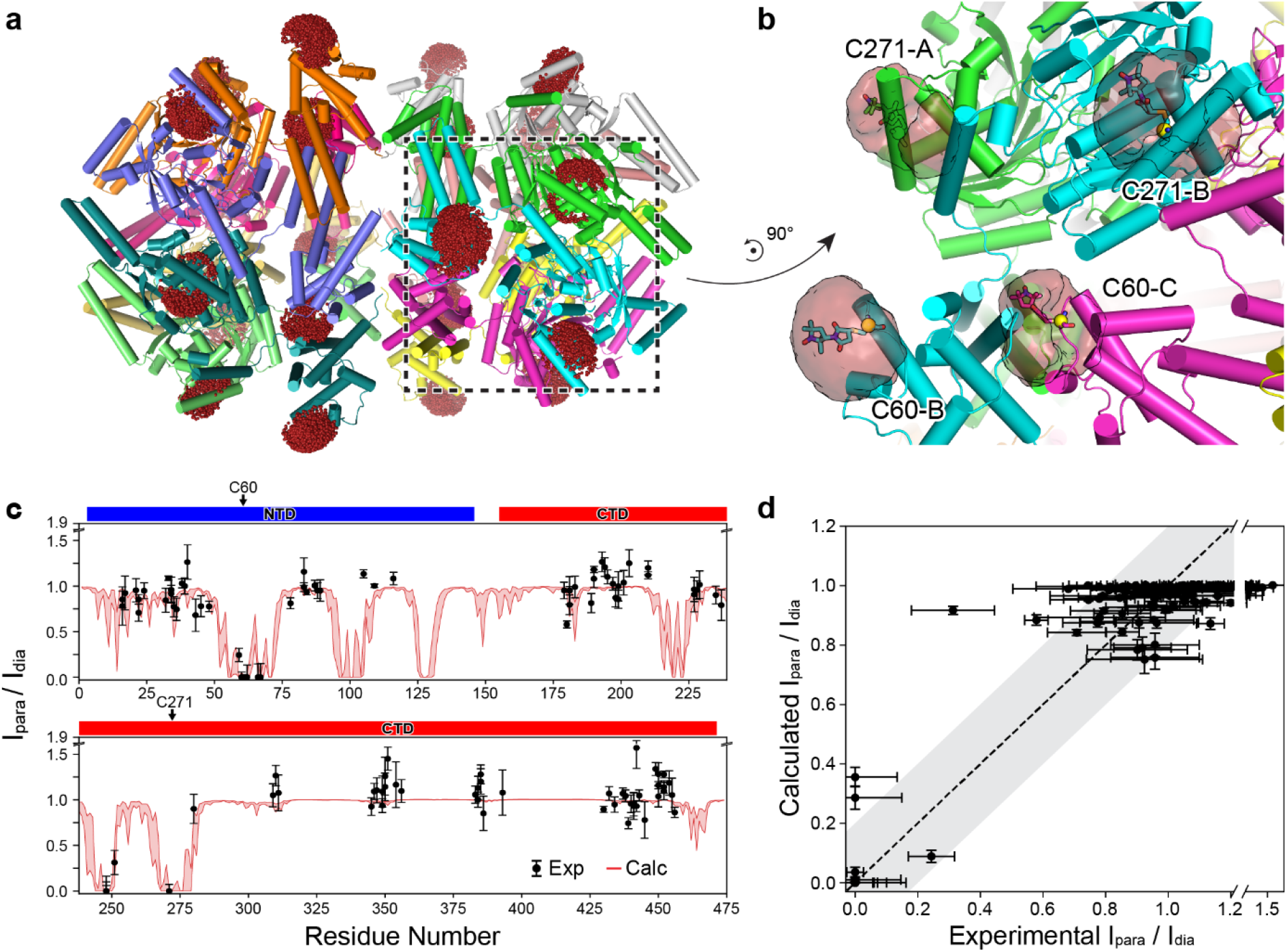
Model-predicted nitroxide spin density of PROXYL-M tags in DnaB dodecamer and comparison between experimental and calculated PREs. (**a**) Side view of DnaB dodecamer in cartoon representation. Helix regions are depicted as cylinders and the O1 atom positions, resulting from modeled conformations of the PROXYL-M tag, are shown as red spheres. (**b**) Zoom in view in DnaB showing four different spin labeling sites located on three adjacent DnaB monomers. O1 atom densities corresponding to distributed conformation models of PROXYL-M are shown as red transparent volumes. For every spin label site, a representative conformer is depicted in sticks. (**c**) Experimentally observed and model-calculated I_para_/I_dia_ signal intensity ratios versus DnaB residue number. The paramagnetic spectrum (I_para_) was measured on PROXYL-M-tagged DnaB and the diamagnetic spectrum (I_dia_) was measured on reduced PROXYL-M-tagged DnaB. Vertical bars correspond to estimated errors for experimental and calculated PREs. The latter represent one standard deviation of 100 trials of a Monte Carlo error estimation protocol in which 50% of the model-predicted spin label-protein residue distances were randomly deleted. **(d**) Experimental vs. calculated I_para_/I_dia_ correlation plot. The gray shaded area corresponds to an uncertainty of I_para_/I_dia_ of ±0.2.

While the determination of PREs in 2D spectra of DnaB is difficult due to the large protein size, we have recently described for DnaB that this problem can, in principle, be mitigated by increasing the number of spectral dimensions[20]. Thus, we also recorded 3D NCACB spectra of PROXYL-M-tagged DnaB and apo DnaB, respectively. Figure S5a displays experimental site-specific PRE data (black data points) and simulated PRE data (red curve) for the 3D NCACB experiment. While the number of site-specifically assigned PRE data is increased in the 3D NCACB spectrum compared to the 2D DARR spectrum, the agreement between experimental and calculated PREs is not as good as for the DARR experiment (for only 56% of PREs the difference was within a margin of +/- 0.2). This could be attributed to the lower signal-to-noise ratios of the 3D spectra leading to larger errors of the experimental PRE values and/or to the high sensitivity of the DREAM transfer step for the chosen experimental conditions. Therefore, we interpreted those PREs only qualitatively in previous studies[21].

Figures 6a and 6b show the positions of the Gd^3+^ ion of Gd^3+^-DOTA-M in the DnaB dodecamer and the Gd^3+^ density at four different spin-label sites in one DnaB homodimer, respectively. While the overall shape of the Gd^3+^ density distribution is similar to that one observed for the nitroxide radical, the space sampled by the Gd^3+^ ion in the computational simulation is normally larger compared to the nitroxide radical. This is likely a consequence of the increased number of side-chain degrees of freedom of Gd^3+^-DOTA-M (seven dihedral angles) versus PROXYL-M (four dihedral angles). Interestingly, the spaces occupied by PROXYL-M and Gd^3+^-DOTA-M at one of the two C60 residues in the DnaB homodimer (labeled C60-C in Figures 5b and 6b) are slightly displaced relative to each other. This could be explained by the fact that C60 is located at the interface between two DnaB NTD dimers, which limits the conformational space for the larger Gd^3+^-DOTA-M tag but leaves more room for the smaller PROXYL-M tag.

**Figure 6:**
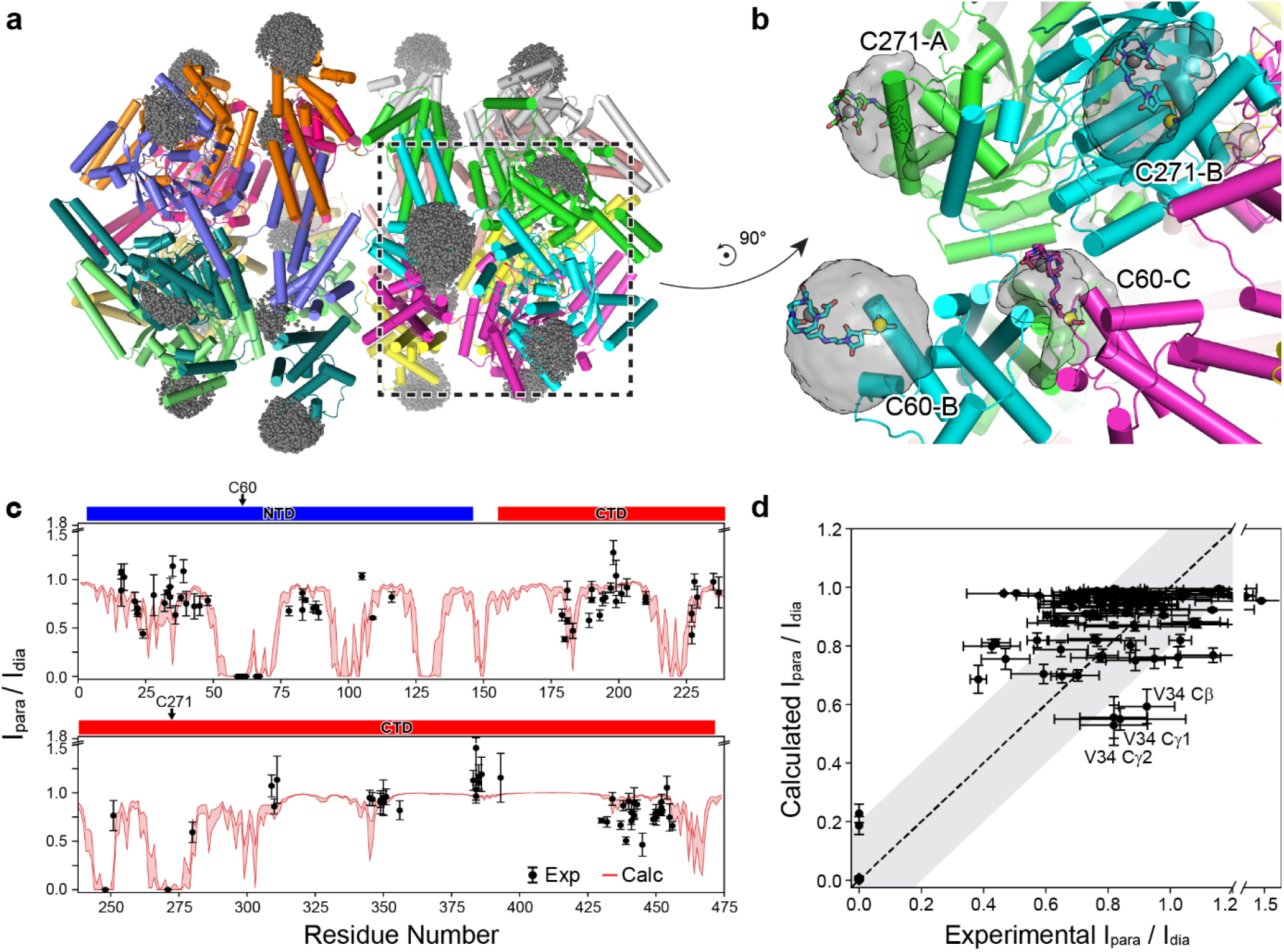
Model-predicted Gd^3+^ density of Gd^3+^-DOTA-M tags in DnaB dodecamer and comparison between experimental and calculated PREs. (**a**) Side view of DnaB dodecamer in cartoon representation. Helix regions are depicted as cylinders and the Gd^3+^ positions of the Gd^3+^-DOTA-M tag are shown as gray spheres. (**b**) Zoom in view in DnaB showing four different spin labeling sites located on three adjacent DnaB monomers. Gd^3+^ ion densities corresponding to ensemble conformation models of Gd^3+^-DOTA-M are shown as gray volumes. For every spin label site, a representative conformer is depicted in sticks. (**c**) Experimentally observed and model-calculated I_para_/I_dia_ signal intensity ratios versus DnaB residue number. The paramagnetic spectrum (I_para_) was measured on DOTA-M-tagged DnaB loaded with Gd^3+^ and the diamagnetic spectrum (I_dia_) was recorded after replacing the metal ion with Lu^3+^. Vertical bars correspond to estimated errors for experimental and calculated PREs. The latter represent one standard deviation of 100 trials of a Monte Carlo error estimation protocol in which 50% of the model-predicted spin label-protein residue distances were randomly deleted. (**d**) Experimental vs. calculated I_para_/I_dia_ correlation plot. The gray shaded area corresponds to an uncertainty of I_para_/I_dia_ of ±0.2.

Figures 6c and 6d compare experimental PREs obtained on Gd^3+^-DOTA-M-tagged DnaB with PREs calculated from the DnaB structure by determining effective Gd^3+^-amino acid residue distances. Overall, structural modeling supports the experimental PRE measurements; for 63% of PREs, the agreement between experimental and calculated values is equal or better ±0.2. Noteworthy, I_para_/I_dia_ values with deviations from the calculated curve clustered around residue positions 400 and 450 in Figure 6c. This could possibly indicate flexibility of the CTD which also escaped detection by solid-state NMR. Another small but noticeable difference between experimentally observed and calculated I_para_/I_dia_ values was found for the cross peaks of V34. This could indicate increased conformational flexibility of the helix α1-α2-connecting loop containing V34. Alternatively, this could indicate some flaws of the X-ray structure model in this loop region, which is poorly defined by the previously reported low-resolution electron density (see for a further discussion the supporting material and Figure S6).

### Determination of the blind sphere of Gd^3+^-DOTA-M and PROXYL-M spin labels

An important consideration for NMR experiments employing paramagnetic spin labels is the size of the so-called “blind sphere”, i.e., the region where a nucleus is sufficiently close to the unpaired electron so that its NMR signal is broadened entirely beyond detection. To determine the blind spheres induced by the nitroxide radical and Gd^3+^ ion for direct detection of ^13^C, we plotted the experimental PRE values versus the model-determined effective nitroxide-carbon and Gd^3+^-carbon distances, respectively, and analyzed the resultant curves by calculating the least-squares fits to the theoretically-derived equation described in reference [18]. Figure 7 and Figures S7 and S8 illustrate how the PRE (I_para_/I_dia_) changes with the nitroxide-carbon or Gd^3+^-carbon distance in the 2D DARR and 3D NCACB experiment, respectively.

**Figure 7:**
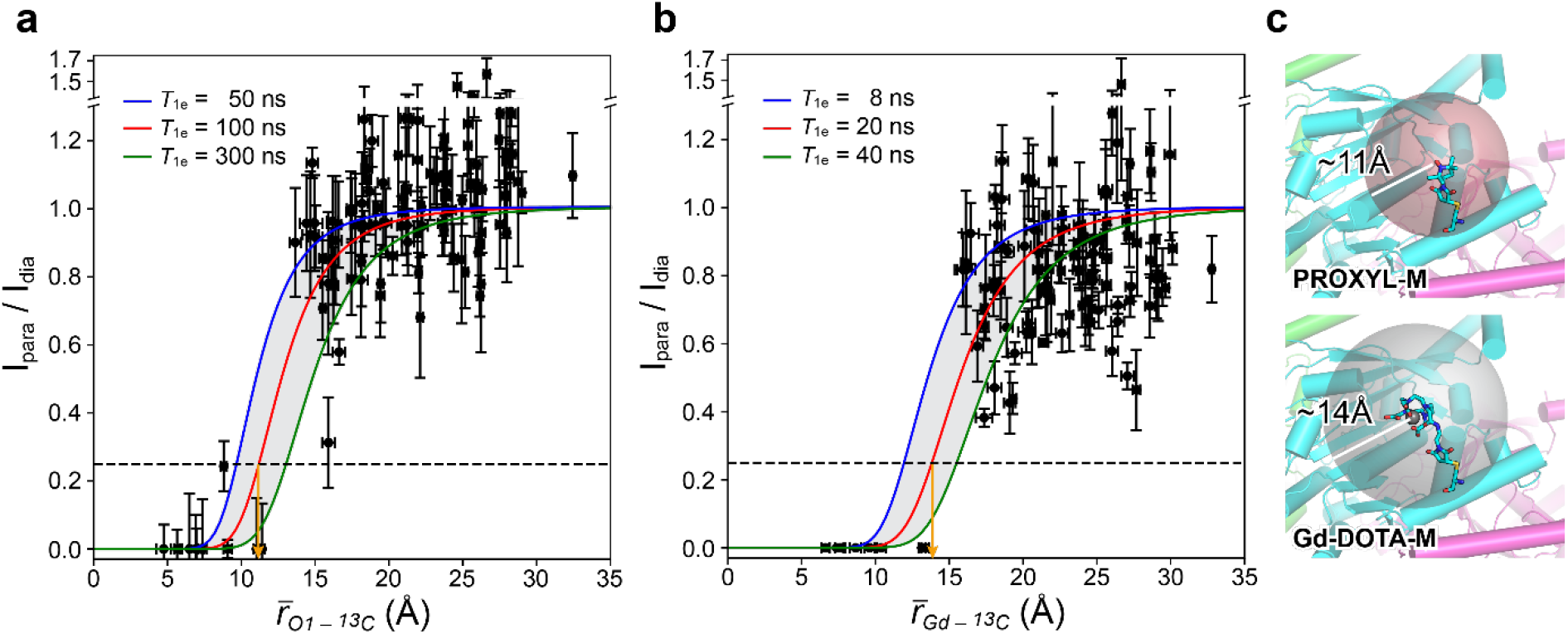
Intensity-vs-spin label distance curves for Gd^3+^-DOTA-M and PROXYL-M. (**a**) I_para_/I_dia_ ratio extracted from 2D spectra versus effective nitroxide-nuclear spin distance for PROXYL-M, and (**b**) I_para_/I_dia_ ratio versus effective Gd^3+^-nuclear spin distance for Gd^3+^-DOTA-M. The dashed line indicates the lower limit of the signal intensity ratio below which the signal in the paramagnetic spectrum is usually too weak to be detected. The colored lines represent least-squares fits to the theoretically-derived equation described in reference [18] assuming different electronic *T*_1e_ relaxation times. A 25% intensity ratio corresponds to an effective distance of 11 Å or 14 Å for PROXYL-M and Gd^3+^-DOTA-M tags, respectively, which marks the size of the blind sphere for these tags (orange arrows). (**c**) Visual comparison of the size of the blind sphere for PROXYL-M and Gd-DOTA-M tags in DnaB.

Best fits were obtained with *T*_1e_-values of 100 ns for the nitroxide radical and 20 ns for the Gd^3+^ ion, which agrees well with previously determined values for *T*_1e_[60]. Assuming a lower detection limit for I_para_/I_dia_ of 25%, below which an NMR signal is usually too weak to be detected in the paramagnetic spectrum, these fits yielded estimates for the size of the ^13^C blind sphere in 2D DARR spectra of 11 Å and 14 Å for the nitroxide and Gd^3+^, respectively. In comparison, the nitroxide tag leads to a slightly larger ^13^C blind sphere in the 3D NCACB spectrum of 12 Å (Figure S5c and S8). These values compare favorably with the reported ones for the nitroxide in MTSL[61] or the Gd^3+^ ion in dopants and metalloproteins[62, 63], as well as in metal-ion DNP studies employing Gd^3+^-DOTA-M tags[26]. Noteworthy, the lengths of the linker connecting the protein backbone to the nitroxide (10.2 Å) or the Gd^3+^ ion (12.5 Å) in the model structures is only slightly smaller than the size of the blind sphere determined in Figure 7. A sufficiently large linker ensures that the unpaired electron is removed from the protein surface, and, thus, avoids that too many residues get within the blind region of the spin label, which is relevant for instance in protein-protein interaction studies. If, however, the blind sphere information shall be used in structure determination, tags with shorter linkers are even favored.

### EPR distance measurements using Gd^3+^-DOTA-M-taggedDnaB

Spin labeled DnaB also opens the way for EPR distance measurements using dipolar electron-electron resonance (DEER) spectroscopy[64-66]. We recorded Gd^3+^-Gd^3+^ DEER[67] data after labeling DnaB with Gd^3+^-DOTA-M at a single cysteine site. To this end, residue T116 was replaced by a cysteine and the two native cysteine residues, C60 and C271, were mutated to an alanine and a serine, respectively. The DEER distance distribution in Figure 8a shows three maxima at around 3.8, 6, and 7.5 nm, although the latter cannot be extracted with high confidence. We compared the location of the maxima in the experimental distance distribution to the ones in the distance distribution obtained by molecular modeling with Rosetta (Figure 8b) and found a good overall agreement. Such a distance distribution is consistent with a labeling of all available T116C sites in the DnaB crystal structure model (Figure 8c) and therefore serves as a spectroscopic evidence that DnaB was present as a dimer of hexamers (or dodecameric assembly). Similar experiments will also allow us in the future to probe interactions of DnaB with other proteins, such as with the DnaG primase within the primosome complex.

**Figure 8:**
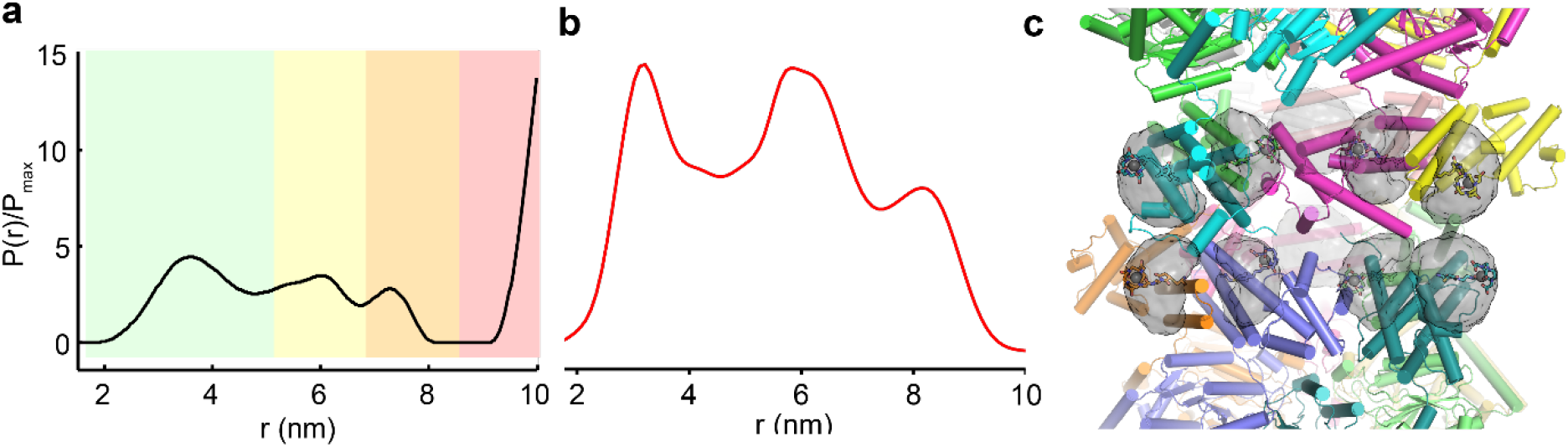
Gd^3+^-Gd^3+^DEER distance distribution obtained on Gd^3+^-DOTA-M-tagged DnaB is consistent with a hexameric subunit stoichiometry. (a) Experimental Gd^3+^-Gd^3+^ DEER distance distribution at 10 K, green: shape of distance distribution is reliable; yellow: mean distance and width are reliable; orange: mean distance is reliable; red: long-range distance contributions may be detectable, but cannot be quantified. Theoretically predicted distance distributions using Rosetta (b). (c) Gd^3+^ spin densities on the lanthanide-tagged DnaB mutant C60A/T116C/C271A as calculated with Rosetta.

## Conclusions

We herein describe the site-directed covalent spin labeling of the bacterial DnaB helicase from *Helicobacter pylori* with PROXYL-M and DOTA-M tags. DnaB constitutes a large protein with 488 residues per monomer and we show that paramagnetic solid-state NMR on this oligomeric assembly becomes possible opening the way for structural investigations. Site-specific PRE data were extracted from 2D and 3D solid-state NMR spectra and agreed well with PREs back-calculated from the DnaB structure by computational modeling using the low-resolution DnaB crystal structure. The radius of the blind sphere of PROXYL-M in ^13^C-^13^C DARR experiments (~11 Å) is smaller than that of Gd^3+^-DOTA-M (~14 Å). This information will be of practical relevance for the design of future NMR studies using these paramagnetic tags. Our study showcases the combined application of paramagnetic spin labeling, solid-state NMR, and computational modeling for the structural characterization of the large DnaB complex. Integrative technologies like these have the potential to facilitate the structure determination of biomolecular systems that are difficult to study by only a single technique, such as flexible and dynamic macromolecular assemblies and membrane-associated complexes. We also envision that EPR experiments on spin labeled DnaB will be a useful tool providing new insights into the mode of action of DnaB and for large motor proteins in general.

## MATERIALS AND METHODS

### Sample preparation

^13^C and ^15^N labeled DnaB was expressed as described in reference [68]. All protein solutions were sedimented[69-71] in the MAS-NMR rotor (16 h at 4 °C at 210,000 x g) using home-built tools[72].

#### Tagging of DnaB with PROXYL-M

DnaB solution (20 mg dissolved in 5 mL buffer (2.5 mM sodium phosphate, pH 7.5, 130 mM NaCl)) was transferred to new buffer (10 mM sodium phosphate, 150 mM NaCl, 10% glycerine, pH 7.5) and desalted with PD-10 column. The protein was concentrated to 40 mg/mL by centrifugation using Millipore concentrator (30 kDa cut-off). The spin-labeling reaction with PROXYL-M was performed at 4 °C and pH 7.5 for 18 h using a protein concentration of 50 μM and a 30-fold excess of spin label (per cysteine). Finally, the solution was desalted with PD-10 column.

#### Tagging of DnaB with the reduced form of PROXYL-M

DnaB solution (32 mg dissolved in 10 mL buffer (2.5 mM sodium phosphate, pH 7.5, 130 mM NaCl)) was transferred to new buffer (10 mM sodium phosphate, 150 mM NaCl, 10% glycerine, pH 7.5) and desalted with PD-10 column. The protein was concentrated to 7 mg/mL by centrifugation using Millipore concentrator (30 kDa cut-off). The spin labeling rection with PROXYL-M was performed at 4 °C and pH 7.5 for 18 h using a protein concentration of 110 μM and a 30-fold excess of spin label (per cysteine). The reduction of the PROXYL-M tag was performed in a new buffer (10 mM sodium phosphate, 150 mM NaCl, 30 mM ascorbic acid, 10% glycerine, pH 7.5)[22]. Finally, the solution was desalted with PD-10 column.

#### Tagging of DnaB with Lu^3+^-DOTA-M

DnaB solution (35 mg dissolved in 9 mL buffer (2.5 mM sodium phosphate, pH 7.5, 130 mM NaCl)) was transferred to new buffer (10 mM sodium phosphate, 150 mM NaCl, 10% glycerine, pH 7.5) and desalted with PD-10 column. The protein was concentrated to 15 mg/mL by centrifugation using Millipore concentrator (30 kDa cut-off). The spin labeling reaction with Lu^3+^-DOTA-M was performed at 4 °C and pH 7.5 for 18 h using a protein concentration of 160 μM and a 30-fold excess of spin label (per cysteine). Finally, the solution was desalted with PD-10 column.

#### Tagging of DnaB with Gd^3+^-DOTA-M

DnaB solution (35 mg dissolved in 9 mL buffer (2.5 mM sodium phosphate, pH 7.5, 130 mM NaCl)) was transferred to new buffer (10 mM sodium phosphate, 150 mM NaCl, 10% glycerine, pH 7.5) and desalted with PD-10 column. The protein was concentrated to 15 mg/mL by centrifugation using Millipore concentrator (30 kDa cut-off). The spin labeling reaction with Gd^3+^-DOTA-M was performed at 4 °C and pH 7.5 for 18 h using a protein concentration of 160 μM and a 30-fold excess of spin label (per cysteine). Finally, the solution was desalted with PD-10 column.

#### EPR sample preparation

##### DnaB with PROXYL-M

DnaB with PROXYL-M solution (1.09 mg dissolved in 25 μL buffer (10 mM sodium phosphate, 150 mM NaCl, 10% glycerine, pH 7.5)) was used for the CW-EPR experiment.

##### PROXLY-M reference solution

A 3.4 mM PROXYL-M stock solution in buffer (10 mM sodium phosphate, 150 mM NaCl, 10% glycerine, pH 7.5) was prepared. The solution was diluted with buffer (10 mM sodium phosphate, 150 mM NaCl, 10% glycerine, pH 7.5) until a concentration of 100 μM PROXYL-M was reached.

##### DnaB-C60A C271S T116C with Gd^3+^-DOTA-M

DnaB with Gd^3+^-DOTA-M solution (0.25 mg dissolved in 0.5 mL buffer (10 mM sodium phosphate, pH 7.5, 150 mM NaCl)) was transferred to new buffer (10 mM sodium phosphate, 150 mM NaCl, 10% glycerine, pH 7.5) and concentrated to 2.5 mg/mL by centrifugation using Millipore concentrator (10 kDa cut-off). DnaB solution was transferred to new buffer (10 mM sodium phosphate, 150 mM NaCl, pH 7.8, D_2_O) and concentrated to 3.6 mg/mL by centrifugation using Millipore concentrator (10 kDa cut-off). The mutant C60A C271S T116 was expressed as described above for the wild-type protein.

### Solid-state NMR

Solid-state NMR spectra were acquired on a wide-bore 850 MHz Bruker Avance III spectrometer using a 3.2mm Bruker Biospin “E-free” probe[73]. The MAS frequency was set to 17.0 kHz and the sample temperature was adjusted to 278 K using the water line as an internal reference[72]. The 2D spectra were processed with the software TOPSPIN (version 3.5, Bruker Biospin) with a shifted (3.0) squared cosine apodization function and automated baseline correction in the indirect and direct dimension. Spectral analysis was performed using CcpNmr Analysis 2.4.2 [74-76]. The spectra were referenced to 4,4-dimethyl-4-silapentane-1-sulfonic acid (DSS). For all experimental details see Table S4.

### EPR

#### Continuous-wave EPR spectroscopy

CW EPR spectra of nitroxide spin-labelled protein were recorded at room temperature at X band (9.5 GHz) using a Bruker ElexsysE500 spectrometer including a Bruker super high Q resonatorER4122SHQ. Protein samples were filled into 50 μL glass capillaries with an outer/inner diameter of 1.5/0.9 mm (BLAUBRANDs). All spectra were recorded with 100 kHz field modulation, 1 G modulation amplitude, a time constant of 10.24 ms, a conversion time of 40.96 ms and 25 dB microwave power attenuation. Spin labeling efficiencies were calculated by digital double integration of the recorded spectra and comparison with the reference EPR measurement of nitroxide solution of 100 μM concentration.

#### DEER spectroscopy

Pulsed EPR measurements were performed at a home-built high power Q-band spectrometer (35 GHz) with 200 W output micro-wave power, a Bruker ElexSys acquisition system (E580) and a home-built TE001 pulse probe (OD 3 mm sample tubes). Temperature stabilisation during all measurements was performed with a He-flow cryostat (ER 4118CF, Oxford Instruments) and a temperature control system (ITC 503, Oxford Instruments). The four pulse DEER experiment was used to measure Gd(III)-Gd(III) distances at 10 K. The frequency offset between detection and pump pulse was 100 MHz, all pulse lengths were set to 12 ns, the primary echo delay time was set to 400 ns, with pump pulse step of 12 ns and an eight-step ESEEM averaging cycle with a step of 8 ns. Intermolecular couplings, which cause background decay in the time domain data, were removed by background correction (with dimension parameter D=3.0) and distance distributions in a range of 1.5 to 10.0 nm were obtained by Tikhonov regularization.

### Structural modeling of Gd-DOTA-M and PROXYL-M tags in DnaB

The structures of Gd-DOTA-M and PROXYL-M tags conjugated to the side-chain of cysteine (termed Gd-DOTA-M-Cys or PROXYL-M-Cys, respectively) were built in Avogadro (version1.2)[77]. Gd-DOTA-M-Cys was built in two absolute configurations (R and S) to account for the presence of a chiral carbon atom in the Cys-linked maleimide ring. For PROXYL-M-Cys, the structures of four stereoisomers were considered in molecular modeling because of two chiral atoms, one at the maleimide ring and another at the PROXYL ring (see Figure S4). Structure geometry was optimized by DFT calculations with Gaussian09 (Gaussian, Inc., Wallingford CT) using the quasi-relativistic effective core potential (ECP)[78] and [5s4p3d]-GTO valence basis sets for the Gd^3+^ ion, the 6-31(d,p) standard basis set for all other atoms, and the M06 functional[79]. Solvation effects were evaluated with the integral equation formalism of the polarizable continuum model (IEFPCM)[80] implemented in Gaussian09.

The BCL::ConformerGenerator method[81] was used to build libraries of 2000 unique conformations for each stereoisomer of Gd-DOTA-M-Cys and PROXYL-M-Cys, respectively. The method generates 3D ligand conformers by combining rotamers of small molecule structures from the Crystallography Open Database (COD)[58, 59] according to a statistically-derived scoring function. For Gd-DOTA-M-Cys, the linker connecting the DOTA chelator to the backbone Cα atom was fully flexible in conformer generation, while the Gd-DOTA complex was treated as rigid. For PROXYL-M-Cys, all four side-chain dihedral angles were flexible. A total of 20000 conformer generation iterations were carried out from which the 2000 best-scoring conformations were kept after removing similar conformers with a pairwise root-mean-squared distance deviation (RMSD) <0.25Å. The resulting conformer libraries were deemed nearly complete as all of the expected rotamers of the six (Gd-DOTA-M-Cys) or four (PROXYL-M-Cys) dihedral angles occurred with similar probabilities (see Figure S4).

The Rosetta software (version 3.12)[55] was used to model the side-chain conformations and determine the spin density for Gd-DOTA-M and PROXYL-M at every cysteine residue in DnaB. First, the structure of dodecameric DnaB from *H. pylori* (PDB: 4ZC0)[54] was refined by adding missing residues, which were not resolved in the original X-ray structure model, and performing constrained minimization in the Rosetta force field[82]. Modeling of unresolved regions was accomplished by insertion of backbone fragments with PSIPRED-[83] and JUFO9D-[84] predicted secondary structures in combination with cyclic coordinate descent (CCD)[85] loop building algorithms implemented through the Rosetta comparative modeling (RosettaCM)[86] program. In this step, D3 symmetry of DnaB dodecamer was enforced with the help of symmetry definition files[87], and Cα-atom pair distance constraints were applied to prevent any large movements of the protein backbone during loop modeling and minimization. Second, all of the 24 native cysteine residues in DnaB were successively replaced by every conformer of the S- and R-isomers of Gd-DOTA-M-Cys or PROXYL-M-Cys, respectively, by aligning the corresponding backbone atoms of protein and spin label residue with each other. The backbone and side-chain degrees of freedom of all protein residues surrounding the spin label were minimized using the Rosetta all-atom ref2015 energy function[82] while applying weak distance constraints between all pairs of Cα-atoms which were within 10 Å of each other. Spin label conformers which still clashed with the protein after minimization according to a Rosetta energy cutoff (−5000 kcal/mol) were removed. The remaining conformers were used to calculate the spin density distribution for the Gd^3+^ ion or the nitroxide radical, respectively. The molecular surface area of cysteine residues in the DnaB structure model was measured with the *get_area* function in PyMOL (The PyMOL Molecular Graphis System, Version 2.4, Schrödinger, LCC).

### Simulation of PREs for Gd^3+^ and nitroxide spin labels

The intensity ratio of an NMR signal recorded in the para- vs. diamagnetic spectrum was calculated according to:

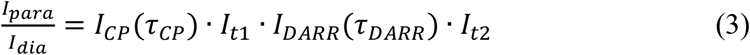

where I_CP_ is the change in signal intensity during the cross polarization (CP) step, I_DARR_ is the signal reduction during the DARR mixing period, and I_t1_ and I_t2_ represent the signal intensity reduction during the evolution (t_1_) and detection (t_2_) periods, respectively. The derivation of equation (1) is described in detail in Tamaki et al. [18]. I_CP_ is dependent on the paramagnetic relaxation rates of ^1^H and ^13^C in the rotating frame (Γ _*1*ρ,H_, Γ _*1*ρ,c_), I_DARR_ is a function of the paramagnetic longitudinal relaxation rate of ^13^C (Γ_1,c_), and I_t1_ and I_t2_ are determined by the paramagnetic transversal relaxation rate of ^13^C (Γ_2,c_). All equations and simulation parameters are provided in the supporting material (see Figures S7-S9).

The effective unpaired electron-nucleus distance, 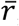, was calculated based on the refined DnaB structure model and the simulated spin density distributions for Gd^3+^ and the nitroxide radical according to[26]:

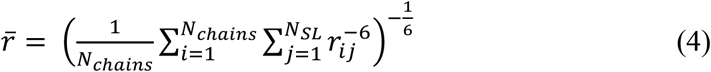

*N*_SL_, is the number of spin label sites (*N*_SL_ = 24) in the DnaB dodecamer, and *r*_ij_ is the distance between a carbon atom in DnaB, for which a cross peak was detected in the NMR experiment, and the center of mass of the Gd^3+^ ion (DOTA-M) or O1 atom (PROXYL-M) positions, respectively, for each of the 24 spin label sites. In calculating the center of mass of the spin density, we assumed an N-jump model for the spin label in accordance with previous approaches[57, 88]. Because the dipolar relaxation of a nucleus in one DnaB monomer is the combined effect of all 24 spin labels in DnaB dodecamer, the effective distance in equation (4) is evaluated by the sum of r^-6^ distances to the Gd^3+^ or O1 centers of mass of all 24 spin label sites. In addition, the distances to both DnaB carbon atoms, which contribute to the same NMR cross peak, are included in the summation in equation (4). Note that because the conformations of chains A and B in the DnaB crystal structure are not identical (Cα-RMSD is 4.6 Å over 72% of the residues), r^-6^ averaging of the distances calculated for chains A and B is additionally applied in equation (4). Thus, *N*_chains_ denotes the number of asymmetric protein chains (A and B, *N*_chains_ = 2) in the DnaB dodecamer.

## Author contributions

JZ and TW performed the NMR experiments, RC prepared the samples. MY recorded and analyzed the EPR spectra. GK performed the computational modeling. JZ, MY, GK and TW analyzed the data. GK and TW designed the research. JZ, GK and TW wrote the initial manuscript draft. All authors reviewed and edited the manuscript.

## Acknowledgements

This work was supported by the ETH Research Grant ETH-43 17-2 (T.W.) and the Deutsche Forschungsgemeinschaft (DFG, German Research Foundation, project number 455240421 and Heisenberg fellowship, T.W.). GK was supported by a DFG postdoctoral fellowship (KU 3510/1-1). TW acknowledges helpful discussions with Prof. Beat H. Meier (ETH Zürich, Switzerland), Dr. Anja Böckmann and Dr. Laurent Terradot (both MMSB Lyon, France) as well as Dr. Carole Gardiennet (Université de Lorraine, France). Prof. Beat H. Meier is also acknowledged for providing NMR measurement time.

## References

[1] M. Bonaccorsi, T. Le Marchand, G. Pintacuda, Protein structural dynamics by Magic-Angle Spinning NMR, Curr. Opin. Struct. Biol., 70 (2021) 34–43.

[2] P.C.A. van der Wel, New applications of solid-state NMR in structural biology, Emerg. Top. Life Sci., 2 (2018) 57–67.

[3] E. Nimerovsky, K.T. Movellan, X.C. Zhang, M.C. Forster, E. Najbauer, K. Xue, R. Dervişoğlu, K. Giller, C. Griesinger, S. Becker, L.B. Andreas, Proton Detected Solid-State NMR of Membrane Proteins at 28 Tesla (1.2 GHz) and 100 kHz Magic-Angle Spinning, Biomolecules, 11 (2021) 752.

[4] M. Callon, A.A. Malär, S. Pfister, V. Římal, M.E. Weber, T. Wiegand, J. Zehnder, M. Chávez, R. Cadalbert, R. Deb, A. Däpp, M.-L. Fogeron, A. Hunkeler, L. Lecoq, A. Torosyan, D. Zyla, R. Glockshuber, S. Jonas, M. Nassal, M. Ernst, A. Böckmann, B.H. Meier, Biomolecular solid-state NMR spectroscopy at 1200 MHz: the gain in resolution, J. Biomol. NMR, (2021).

[5] A. Böckmann, M. Ernst, B.H. Meier, Spinning proteins, the faster, the better?, J. Magn. Reson., 253 (2015) 71–79.

[6] A. Samoson, H-MAS, J. Magn. Reson., 306 (2019) 167–172.

[7] B. Habenstein, C. Wasmer, L. Bousset, Y. Sourigues, A. Schuetz, A. Loquet, B.H. Meier, R. Melki, A. Böckmann, Extensive de novo solid-state NMR assignments of the 33 kDa C-terminal domain of the Ure2 prion, J. Biomol. NMR, 51 (2011) 235–243.

[8] Y. Shen, F. Delaglio, G. Cornilescu, A. Bax, TALOS+: a hybrid method for predicting protein backbone torsion angles from NMR chemical shifts, J. Biomol. NMR, 44 (2009) 213–223.

[9] G. Pintacuda, G. Kervern, Paramagnetic Solid-State Magic-Angle Spinning NMR Spectroscopy, in: H. Heise, S. Matthews (Eds.) Modern NMR Methodology, Springer Berlin Heidelberg, 2013, pp. 157–200.

[10] G. Otting, Protein NMR Using Paramagnetic Ions, Ann. Rev. Biophys., 39 (2010) 387–405.

[11] G. Otting, Prospects for lanthanides in structural biology by NMR, J. Biomol. NMR, 42 (2008) 1–9.

[12] C.P. Jaroniec, Solid-state nuclear magnetic resonance structural studies of proteins using paramagnetic probes, Solid State Nucl. Magn. Reson., 43-44 (2012) 1–13.

[13] G.M. Clore, J. Iwahara, Theory, Practice, and Applications of Paramagnetic Relaxation Enhancement for the Characterization of Transient Low-Population States of Biological Macromolecules and Their Complexes, Chem. Rev., 109 (2009) 4108–4139.

[14] A.J. Pell, G. Pintacuda, C.P. Grey, Paramagnetic NMR in solution and the solid state, Prog. Nucl. Magn. Reson. Spectrosc., 111 (2019) 1–271.

[15] I. Solomon, Relaxation Processes in a System of Two Spins, Phys. Rev., 99 (1955) 559–565.

[16] N. Bloembergen, L.O. Morgan, Proton Relaxation Times in Paramagnetic Solutions. Effects of Electron Spin Relaxation, J. Chem. Phys., 34 (1961) 842–850.

[17] C.P. Jaroniec, Structural studies of proteins by paramagnetic solid-state NMR spectroscopy, J Magn Reson, 253 (2015) 50–59.

[18] H. Tamaki, A. Egawa, K. Kido, T. Kameda, M. Kamiya, T. Kikukawa, T. Aizawa, T. Fujiwara, M. Demura, Structure determination of uniformly 13C, 15N labeled protein using qualitative distance restraints from MAS solid-state 13C-NMR observed paramagnetic relaxation enhancement, J. Biomol. NMR, 64 (2016) 87–101.

[19] S. Balayssac, I. Bertini, M. Lelli, C. Luchinat, M. Maletta, Paramagnetic Ions Provide Structural Restraints in Solid-State NMR of Proteins, J. Am. Chem. Soc., 129 (2007) 2218–2219.

[20] J. Zehnder, R. Cadalbert, L. Terradot, M. Ernst, A. Bockmann, P. Guntert, B.H. Meier, T. Wiegand, Paramagnetic Solid-State NMR to Localize the Metal-Ion Cofactor in an Oligomeric DnaB Helicase, Chemistry, 27 (2021) 7745–7755.

[21] T. Wiegand, D. Lacabanne, K. Keller, R. Cadalbert, L. Lecoq, M. Yulikov, L. Terradot, G. Jeschke, B.H. Meier, A. Böckmann, Solid-state NMR and EPR Spectroscopy of Mn2+-Substituted ATP-Fueled Protein Engines, Angew. Chem. Int. Ed., 56 (2017) 3369–3373.

[22] M. Ahmed, A. Marchanka, T. Carlomagno, Structure of a Protein-RNA Complex by Solid-State NMR Spectroscopy, Angew. Chem. Int. Ed. Engl., 59 (2020) 6866–6873.

[23] P.S. Nadaud, J.J. Helmus, S.L. Kall, C.P. Jaroniec, Paramagnetic Ions Enable Tuning of Nuclear Relaxation Rates and Provide Long-Range Structural Restraints in Solid-State NMR of Proteins, J. Am. Chem. Soc., 131 (2009) 8108–8120.

[24] M.R. Ermácora, J.M. Delfino, B. Cuenoud, A. Schepartz, R.O. Fox, Conformation-dependent cleavage of staphylococcal nuclease with a disulfide-linked iron chelate, Proc. Natl. Acad. Sci., 89 (1992) 6383–6387.

[25] X.-C. Su, G. Otting, Paramagnetic labelling of proteins and oligonucleotides for NMR, J. Biomol. NMR, 46 (2010) 101–112.

[26] J. Heiliger, T. Matzel, E.C. Çetiner, H. Schwalbe, G. Kuenze, B. Corzilius, Site-specific dynamic nuclear polarization in a Gd(iii)-labeled protein, Phys. Chem. Chem. Phys., 22 (2020) 25455–25466.

[27] S. Wang, R.A. Munro, S.Y. Kim, K.-H. Jung, L.S. Brown, V. Ladizhansky, Paramagnetic Relaxation Enhancement Reveals Oligomerization Interface of a Membrane Protein, J. Am. Chem. Soc., 134 (2012) 16995–16998.

[28] M. Tang, D.A. Berthold, C.M. Rienstra, Solid-State NMR of a Large Membrane Protein by Paramagnetic Relaxation Enhancement, J. Phys. Chem. Lett., 2 (2011) 1836–1841.

[29] E. Teissier, G. Zandomeneghi, A. Loquet, D. Lavillette, J.-P. Lavergne, R. Montserret, F.-L. Cosset, A. Böckmann, B.H. Meier, F. Penin, E.-I. Pécheur, Mechanism of Inhibition of Enveloped Virus Membrane Fusion by the Antiviral Drug Arbidol, PLoS ONE, 6 (2011) e15874.

[30] V. Jirasko, A. Lends, N.-A. Lakomek, M.-L. Fogeron, M.E. Weber, A.A. Malär, S. Penzel, R. Bartenschlager, B.H. Meier, A. Böckmann, Dimer Organization of Membrane-Associated NS5A of Hepatitis C Virus as Determined by Highly Sensitive 1H-Detected Solid-State NMR, Angew. Chem. Int. Ed., 60 (2021) 5339–5347.

[31] X. Jia, H. Yagi, X.-C. Su, M. Stanton-Cook, T. Huber, G. Otting, Engineering [Ln(DPA)3]3-binding sites in proteins: a widely applicable method for tagging proteins with lanthanide ions, J. Biomol. NMR, 50 (2011) 411–420.

[32] Z. Wei, Y. Yang, Q.-F. Li, F. Huang, H.-H. Zuo, X.-C. Su, Noncovalent Tagging Proteins with Paramagnetic Lanthanide Complexes for Protein Study, Chem. Eur. J., 19 (2013) 5758–5764.

[33] R.M. Almeida, C.F.G.C. Geraldes, S.R. Pauleta, J.J.G. Moura, Gd(III) Chelates as NMR Probes of Protein-Protein Interactions. Case Study: Rubredoxin and Cytochrome c3, Inorg. Chem., 50 (2011) 10600–10607.

[34] J.-C. Hus, J.J. Prompers, R. Brüschweiler, Assignment Strategy for Proteins with Known Structure, J. Magn. Reson., 157 (2002) 119–123.

[35] M. Zweckstetter, Determination of molecular alignment tensors without backbone resonance assignment: Aid to rapid analysis of protein-protein interactions, J. Biomol. NMR, 27 (2003) 41–56.

[36] G. Pintacuda, T. Huber, M. Keniry, A. Park, N. Dixon, G. Otting, Fast Assignments of 15N-HSQC Spectra of Proteins by Paramagnetic Labeling, in: G. Webb (Ed.) Modern Magnetic Resonance, Springer Netherlands, 2006, pp. 1263–1269.

[37] P. Shealy, Y. Liu, M. Simin, H. Valafar, Backbone resonance assignment and order tensor estimation using residual dipolar couplings, J. Biomol. NMR, 50 (2011) 357–369.

[38] C. Langmead, B. Donald, An expectation/maximization nuclear vector replacement algorithm for automated NMR resonance assignments, J. Biomol. NMR, 29 (2004) 111–138.

[39] Y.-S. Jung, M. Zweckstetter, Backbone assignment of proteins with known structure using residual dipolar couplings, J. Biomol. NMR, 30 (2004) 25–35.

[40] Y. Tian, C.D. Schwieters, S.J. Opella, F.M. Marassi, AssignFit: A program for simultaneous assignment and structure refinement from solid-state NMR spectra, J. Magn. Reson., 214 (2012) 42–50.

[41] T. Jovanovic, A.E. McDermott, Observation of Ligand Binding to Cytochrome P450 BM-3 by Means of Solid-State NMR Spectroscopy, J. Am. Chem. Soc., 127 (2005) 13816–13821.

[42] S. Balayssac, I. Bertini, A. Bhaumik, M. Lelli, C. Luchinat, Paramagnetic shifts in solid-state NMR of proteins to elicit structural information, Proc. Natl. Acad. Sci., 105 (2008) 17284–17289.

[43] J. Li, K.B. Pilla, Q. Li, Z. Zhang, X. Su, T. Huber, J. Yang, Magic Angle Spinning NMR Structure Determination of Proteins from Pseudocontact Shifts, J. Am. Chem. Soc., 135 (2013) 8294–8303.

[44] I. Bertini, A. Bhaumik, G. De Paëpe, R.G. Griffin, M. Lelli, J.R. Lewandowski, C. Luchinat, High-Resolution Solid-State NMR Structure of a 17.6 kDa Protein, J. Am. Chem. Soc., 132 (2009) 1032–1040.

[45] M.J. Knight, I.C. Felli, R. Pierattelli, I. Bertini, L. Emsley, T. Herrmann, G. Pintacuda, Rapid Measurement of Pseudocontact Shifts in Metalloproteins by Proton-Detected Solid-State NMR Spectroscopy, J. Am. Chem. Soc., 134 (2012) 14730–14733.

[46] P.S. Nadaud, J.J. Helmus, N. Höfer, C.P. Jaroniec, Long-Range Structural Restraints in Spin-Labeled Proteins Probed by Solid-State Nuclear Magnetic Resonance Spectroscopy, J. Am. Chem. Soc., 129 (2007) 7502–7503.

[47] I. Sengupta, P.S. Nadaud, J.J. Helmus, C.D. Schwieters, C.P. Jaroniec, Protein fold determined by paramagnetic magic-angle spinning solid-state NMR spectroscopy, Nature Chem., 4 (2012) 410–417.

[48] I. Sengupta, P.S. Nadaud, C.P. Jaroniec, Protein Structure Determination with Paramagnetic Solid-State NMR Spectroscopy, Acc. Chem. Res., 46 (2013) 2117–2126.

[49] M. Stelter, I. Gutsche, U. Kapp, A. Bazin, G. Bajic, G. Goret, M. Jamin, J. Timmins, L. Terradot, Architecture of a Dodecameric Bacterial Replicative Helicase, Structure, 20 (2012) 554.

[50] T. Wiegand, R. Cadalbert, D. Lacabanne, J. Timmins, L. Terradot, A. Bockmann, B.H. Meier, The conformational changes coupling ATP hydrolysis and translocation in a bacterial DnaB helicase, Nat. Commun., 10 (2019) 31.

[51] M. Spies, DNA Helicases and DNA Motor Proteins, Springer New York, 2012.

[52] K. Takegoshi, S. Nakamura, T. Terao, 13C-1H dipolar-assisted rotational resonance in magic-angle spinning NMR, Chem. Phys. Lett., 344 (2001) 631–637.

[53] K. Takegoshi, S. Nakamura, T. Terao, 13C-13C polarization transfer by resonant interference recoupling under magic-angle spinning in solid-state NMR, Chem. Phys. Lett., 307 (1999) 295–302.

[54] A. Bazin, M.V. Cherrier, I. Gutsche, J. Timmins, L. Terradot, Structure and primase-mediated activation of a bacterial dodecameric replicative helicase, Nucleic Acids Res., 43 (2015) 8564–8576.

[55] J.K. Leman, B.D. Weitzner, S.M. Lewis, J. Adolf-Bryfogle, N. Alam, R.F. Alford, M. Aprahamian, D. Baker, K.A. Barlow, P. Barth, B. Basanta, B.J. Bender, K. Blacklock, J. Bonet, S.E. Boyken, P. Bradley, C. Bystroff, P. Conway, S. Cooper, B.E. Correia, B. Coventry, R. Das, R.M. De Jong, F. DiMaio, L. Dsilva, R. Dunbrack, A.S. Ford, B. Frenz, D.Y. Fu, C. Geniesse, L. Goldschmidt, R. Gowthaman, J.J. Gray, D. Gront, S. Guffy, S. Horowitz, P.-S. Huang, T. Huber, T.M. Jacobs, J.R. Jeliazkov, D.K. Johnson, K. Kappel, J. Karanicolas, H. Khakzad, K.R. Khar, S.D. Khare, F. Khatib, A. Khramushin, I.C. King, R. Kleffner, B. Koepnick, T. Kortemme, G. Kuenze, B. Kuhlman, D. Kuroda, J.W. Labonte, J.K. Lai, G. Lapidoth, A. Leaver-Fay, S. Lindert, T. Linsky, N. London, J.H. Lubin, S. Lyskov, J. Maguire, L. Malmström, E. Marcos, O. Marcu, N.A. Marze, J. Meiler, R. Moretti, V.K. Mulligan, S. Nerli, C. Norn, S. Ó’Conchúir, N. Ollikainen, S. Ovchinnikov, M.S. Pacella, X. Pan, H. Park, R.E. Pavlovicz, M. Pethe, B.G. Pierce, K.B. Pilla, B. Raveh, P.D. Renfrew, S.S.R. Burman, A. Rubenstein, M.F. Sauer, A. Scheck, W. Schief, O. Schueler-Furman, Y. Sedan, A.M. Sevy, N.G. Sgourakis, L. Shi, J.B. Siegel, D.-A. Silva, S. Smith, Y. Song, A. Stein, M. Szegedy, F.D. Teets, S.B. Thyme, R.Y.-R. Wang, A. Watkins, L. Zimmerman, R. Bonneau, Macromolecular modeling and design in Rosetta: recent methods and frameworks, Nature Methods, 17 (2020) 665–680.

[56] D.N. Frick, C.C. Richardson, DNA Primases, Annu. Rev. Biochem, 70 (2001) 39–80.

[57] J. Iwahara, C.D. Schwieters, G.M. Clore, Ensemble Approach for NMR Structure Refinement against 1H Paramagnetic Relaxation Enhancement Data Arising from a Flexible Paramagnetic Group Attached to a Macromolecule, J. Am. Chem. Soc., 126 (2004) 5879–5896.

[58] S. Grazulis, D. Chateigner, R.T. Downs, A.F.T. Yokochi, M. Quiros, L. Lutterotti, E. Manakova, J. Butkus, P. Moeck, A. Le Bail, Crystallography Open Database - an open-access collection of crystal structures, J. Appl. Crystallogr., 42 (2009) 726–729.

[59] S. Gražulis, A. Daškevič, A. Merkys, D. Chateigner, L. Lutterotti, M. Quirós, N.R. Serebryanaya, P. Moeck, R.T. Downs, A. Le Bail, Crystallography Open Database (COD): an open-access collection of crystal structures and platform for world-wide collaboration, Nucleic Acids Res., 40 (2011) D420–D427.

[60] J. Koehler, J. Meiler, Expanding the utility of NMR restraints with paramagnetic compounds: Background and practical aspects, Prog. Nucl. Magn. Reson. Spectrosc., 59 (2011) 360–389.

[61] P. Rovó, K. Grohe, K. Giller, S. Becker, R. Linser, Proton Transverse Relaxation as a Sensitive Probe for Structure Determination in Solid Proteins, ChemPhysChem, 16 (2015) 3791–3796.

[62] W. Li, Q. Zhang, J.J. Joos, P.F. Smet, J. Schmedt auf der Günne, Blind spheres of paramagnetic dopants in solid state NMR, Phys. Chem. Chem. Phys., 21 (2019) 10185–10194.

[63] I. Bertini, C. Luchinat, G. Parigi, E. Ravera, in: I. Bertini, C. Luchinat, G. Parigi, E. Ravera (Eds.) NMR of Paramagnetic Molecules (Second Edition), Elsevier, Boston, 2017, pp. 1–24.

[64] G. Jeschke, Y. Polyhach, Distance measurements on spin-labelled biomacromolecules by pulsed electron paramagnetic resonance, Phys. Chem. Chem. Phys., 9 (2007) 1895–1910.

[65] O. Schiemann, T.F. Prisner, Long-range distance determinations in biomacromolecules by EPR spectroscopy, Quarterly Reviews of Biophysics, 40 (2007) 1–53.

[66] Y.D. Tsvetkov, A.D. Milov, A.G. Maryasov, Russian Chemical Reviews, 77 (2008) 487.

[67] D. Goldfarb, Gd3+ spin labeling for distance measurements by pulse EPR spectroscopy, Phys. Chem. Chem. Phys., 16 (2014) 9685–9699.

[68] T. Wiegand, C. Gardiennet, R. Cadalbert, D. Lacabanne, B. Kunert, L. Terradot, A. Böckmann, B.H. Meier, Variability and conservation of structural domains in divide-and-conquer approaches, J. Biomol. NMR, 65 (2016) 79–86.

[69] I. Bertini, C. Luchinat, G. Parigi, E. Ravera, B. Reif, P. Turano, Solid-state NMR of proteins sedimented by ultracentrifugation, Proc. Natl. Acad. Sci., 108 (2011) 10396–10399.

[70] C. Gardiennet, A.K. Schütz, A. Hunkeler, B. Kunert, L. Terradot, A. Böckmann, B.H. Meier, A Sedimented Sample of a 59 kDa Dodecameric Helicase Yields High-Resolution Solid-State NMR Spectra, Angew. Chem. Int. Ed., 51 (2012) 7855–7858.

[71] T. Wiegand, D. Lacabanne, A. Torosyan, J. Boudet, R. Cadalbert, F.H. Allain, B.H. Meier, A. Bockmann, Sedimentation Yields Long-Term Stable Protein Samples as Shown by Solid-State NMR, Front Mol Biosci, 7 (2020) 17.

[72] A. Böckmann, C. Gardiennet, R. Verel, A. Hunkeler, A. Loquet, G. Pintacuda, L. Emsley, B. Meier, A. Lesage, Characterization of different water pools in solid-state NMR protein samples, J. Biomol. NMR, 45 (2009) 319–327.

[73] P.L. Gor’kov, R. Witter, E.Y. Chekmenev, F. Nozirov, R. Fu, W.W. Brey, Low-E probe for 19F-1H NMR of dilute biological solids, J. Magn. Reson., 189 (2007) 182–189.

[74] R. Fogh, J. Ionides, E. Ulrich, W. Boucher, W. Vranken, J.P. Linge, M. Habeck, W. Rieping, T.N. Bhat, J. Westbrook, K. Henrick, G. Gilliland, H. Berman, J. Thornton, M. Nilges, J. Markley, E. Laue, The CCPN project: an interim report on a data model for the NMR community, Nat Struct Mol Biol, 9 (2002) 416–418.

[75] T. Stevens, R. Fogh, W. Boucher, V. Higman, F. Eisenmenger, B. Bardiaux, B.-J. van Rossum, H. Oschkinat, E. Laue, A software framework for analysing solid-state MAS NMR data, J. Biomol. NMR, 51 (2011) 437–447.

[76] W.F. Vranken, W. Boucher, T.J. Stevens, R.H. Fogh, A. Pajon, M. Llinas, E.L. Ulrich, J.L. Markley, J. Ionides, E.D. Laue, The CCPN data model for NMR spectroscopy: Development of a software pipeline, Proteins: Structure, Function, and Bioinformatics, 59 (2005) 687–696.

[77] M.D. Hanwell, D.E. Curtis, D.C. Lonie, T. Vandermeersch, E. Zurek, G.R. Hutchison, Avogadro: an advanced semantic chemical editor, visualization, and analysis platform, Journal of Cheminformatics, 4 (2012) 17.

[78] M. Dolg, H. Stoll, A. Savin, H. Preuss, Energy-adjusted pseudopotentials for the rare earth elements, Theoret. Chim. Acta, 75 (1989) 173–194.

[79] Y. Zhao, D.G. Truhlar, The M06 suite of density functionals for main group thermochemistry, thermochemical kinetics, noncovalent interactions, excited states, and transition elements: two new functionals and systematic testing of four M06-class functionals and 12 other functionals, Theor. Chem. Acc., 120 (2008) 215–241.

[80] J. Tomasi, B. Mennucci, R. Cammi, Quantum Mechanical Continuum Solvation Models, Chem. Rev., 105 (2005) 2999–3094.

[81] S. Kothiwale, J.L. Mendenhall, J. Meiler, BCL::Conf: small molecule conformational sampling using a knowledge based rotamer library, Journal of Cheminformatics, 7 (2015) 47.

[82] R.F. Alford, A. Leaver-Fay, J.R. Jeliazkov, M.J. O’Meara, F.P. DiMaio, H. Park, M.V. Shapovalov, P.D. Renfrew, V.K. Mulligan, K. Kappel, J.W. Labonte, M.S. Pacella, R. Bonneau, P. Bradley, R.L. Dunbrack, R. Das, D. Baker, B. Kuhlman, T. Kortemme, J.J. Gray, The Rosetta All-Atom Energy Function for Macromolecular Modeling and Design, J. Chem. Theory Comput., 13 (2017) 3031–3048.

[83] D.T. Jones, Protein secondary structure prediction based on position-specific scoring matrices11Edited by G. Von Heijne, J. Mol. Biol., 292 (1999) 195–202.

[84] Leman, Julia Koehler, R. Mueller, M. Karakas, N. Woetzel, J. Meiler, Simultaneous prediction of protein secondary structure and transmembrane spans, Proteins: Structure, Function, and Bioinformatics, 81 (2013) 1127–1140.

[85] A.A. Canutescu, R.L. Dunbrack Jr., Cyclic coordinate descent: A robotics algorithm for protein loop closure, Protein Sci., 12 (2003) 963–972.

[86] Y. Song, F. DiMaio, Ray Y.-R. Wang, D. Kim, C. Miles, T.J. Brunette, J. Thompson, D. Baker, High-Resolution Comparative Modeling with RosettaCM, Structure, 21 (2013) 1735–1742.

[87] F. DiMaio, A. Leaver-Fay, P. Bradley, D. Baker, I. André, Modeling Symmetric Macromolecular Structures in Rosetta3, PLOS ONE, 6 (2011) e20450.

[88] G. Hagelueken, R. Ward, J.H. Naismith, O. Schiemann, MtsslWizard: In Silico Spin-Labeling and Generation of Distance Distributions in PyMOL, Appl. Magn. Reson., 42 (2012) 377–391.

