## Supplementary Material for "Paramagnetic spin labeling of a bacterial DnaB helicase for solid-state NMR"

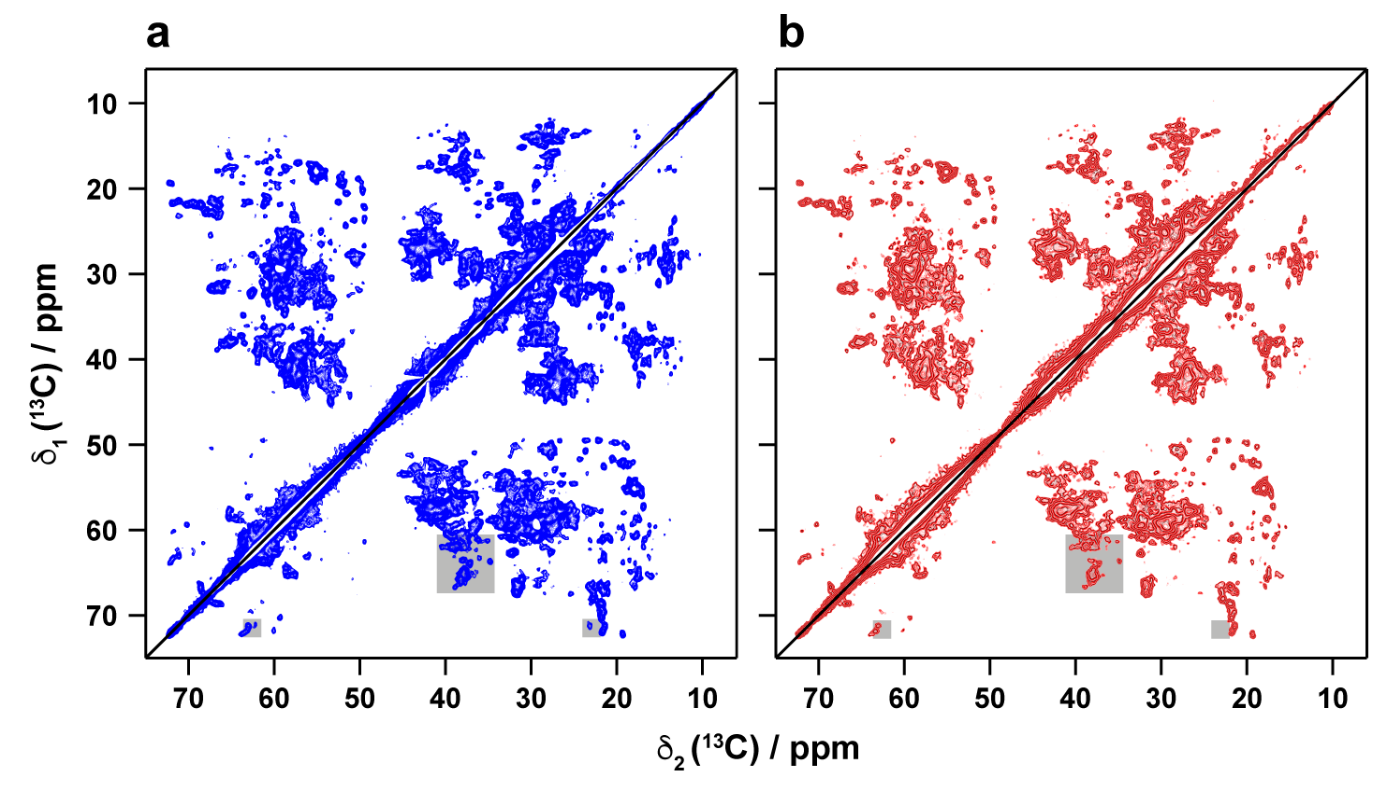

**Figure S1:** 2D ^13^C-^13^C DARR spectra (with 20 ms mixing time) of (**a**) DnaB spin-labeled with PROXYL-M tag and reduced with ascorbic acid (blue) and (**b**) DnaB spin-labeled with PROXYL-M (red). Extracts corresponding to gray regions are shown in Figure 3 in the main text and Figure S2.

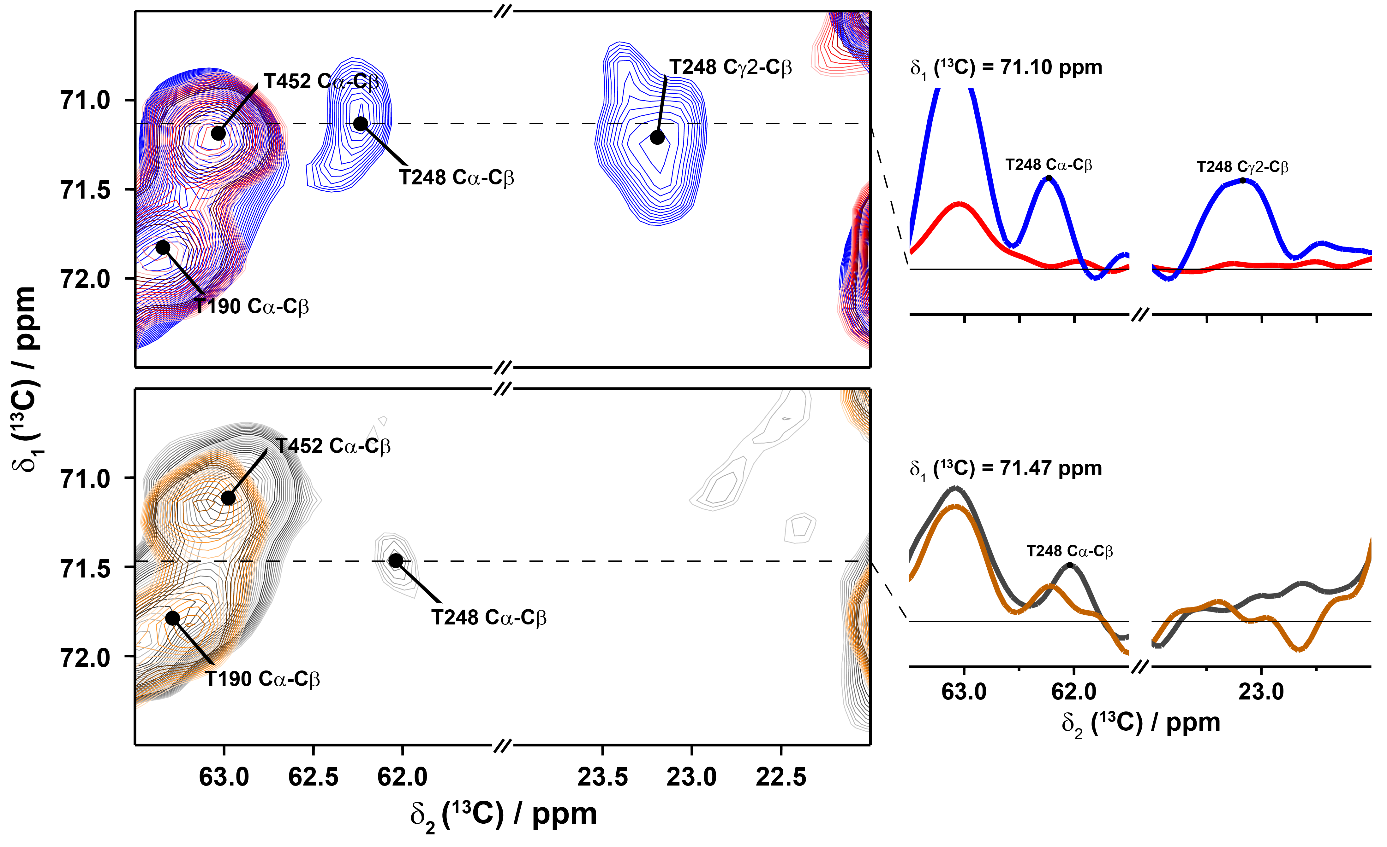

**Figure S2*:*** *Site-specific PRE effects detected on the amino acid T248 in ^13^C-^13^C DARR correlation spectra.* (**top**) Overlay of spectral fingerprints of DnaB spin-labeled with a reduced PROXYL-M tag (blue) and DnaB spin-labeled with PROXYL-M (red). 1D traces along F2 show the cancellation of the resonances corresponding to T248. (**bottom**) Overlay of spectral fingerprints of diamagnetic Lu^3+^-DOTA-M DnaB (gray) and DnaB spin-labeled with Gd^3+^-DOTA-M (orange). 1D traces along F2 show the cancellation of the NMR signal of T248.

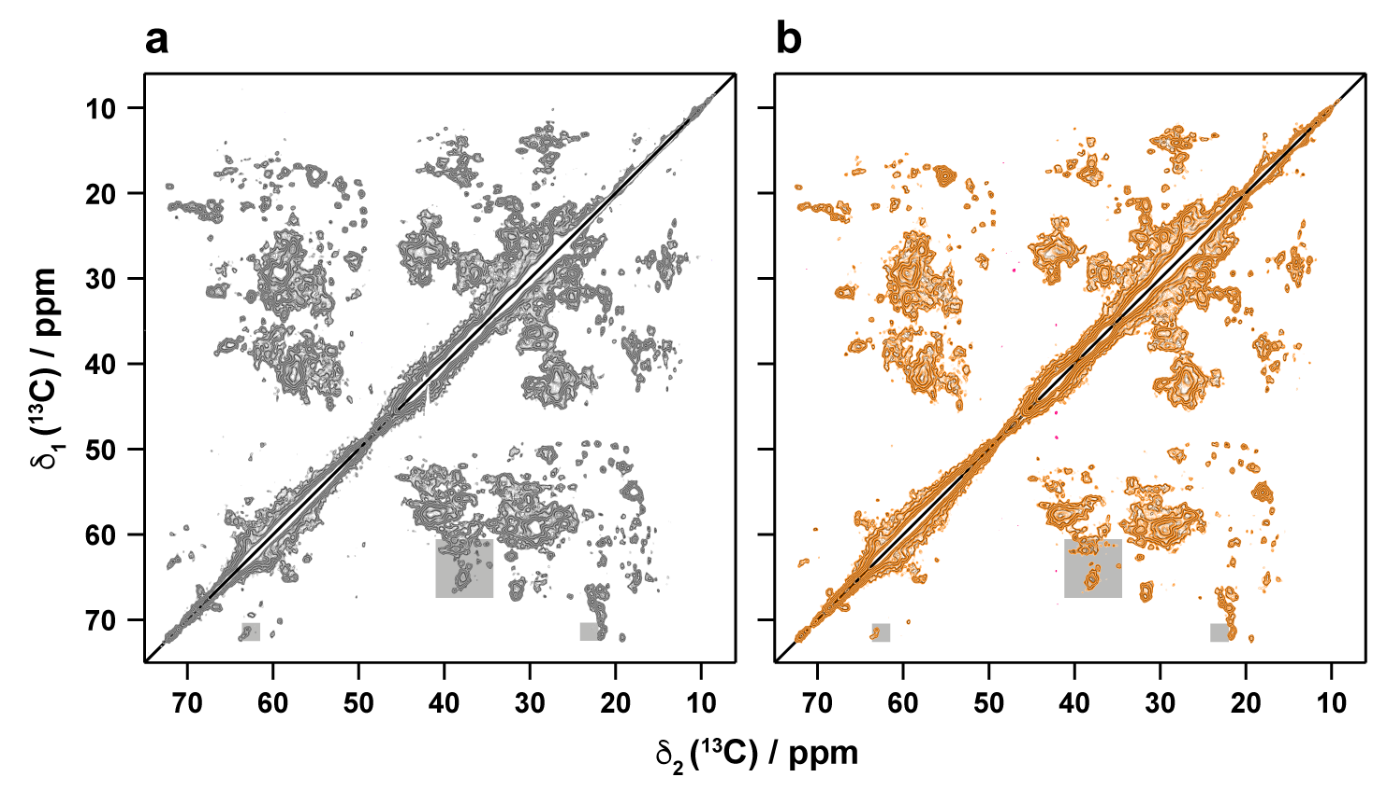

**Figure S3:** 2D ^13^C-^13^C DARR spectra (with 20 ms mixing time) of (**a**) diamagnetic Lu^3+^-DOTA-M DnaB (gray) and (**b**) DnaB spin-labeled with Gd^3+^-DOTA-M (orange). Extracts corresponding to gray regions are shown in Figure 4 in the main text and Figure S2.

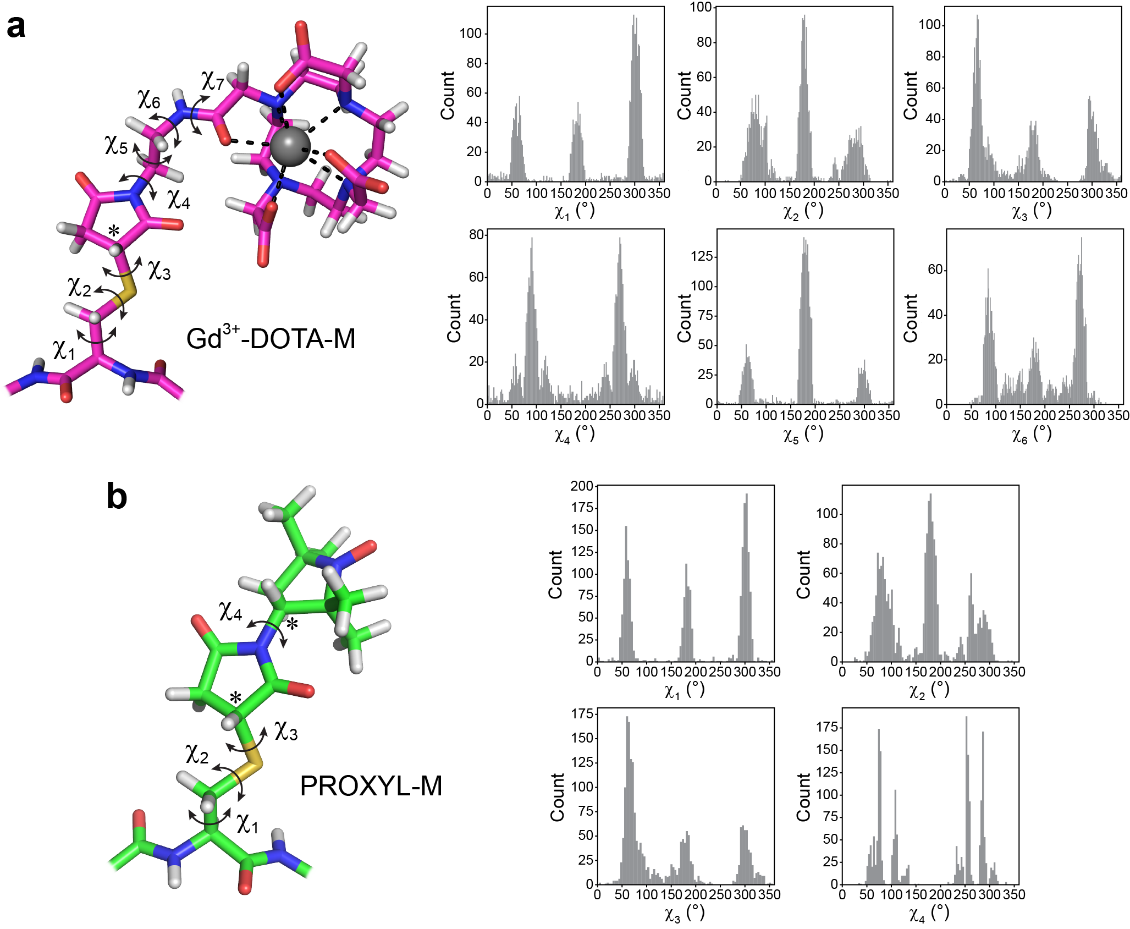

**Figure S4:** *Molecular structures and rotamer libraries of Cys-linked Gd^3+^-DOTA-M and PROXYL-M.* **(a)** Left: Structure of Gd^3+^-DOTA-M-Cys calculated on the M06/[5s4p3d]-GTO/6-31(d,p) level of theory. The side chain dihedral angles and the chiral carbon atom (*) are indicated. Right: Calculated rotamer distributions for χ1 – χ6 angles of Gd^3+^-DOTA-M-Cys. **(b)** Left: Structure of PROXYL-M-Cys calculated on the M06/6-31(d,p) level of theory. The side chain dihedral angles and two chiral carbon atoms (*) are indicated. Right: Calculated rotamer distributions for χ1 – χ4 angles of PROXYL-M-Cys.

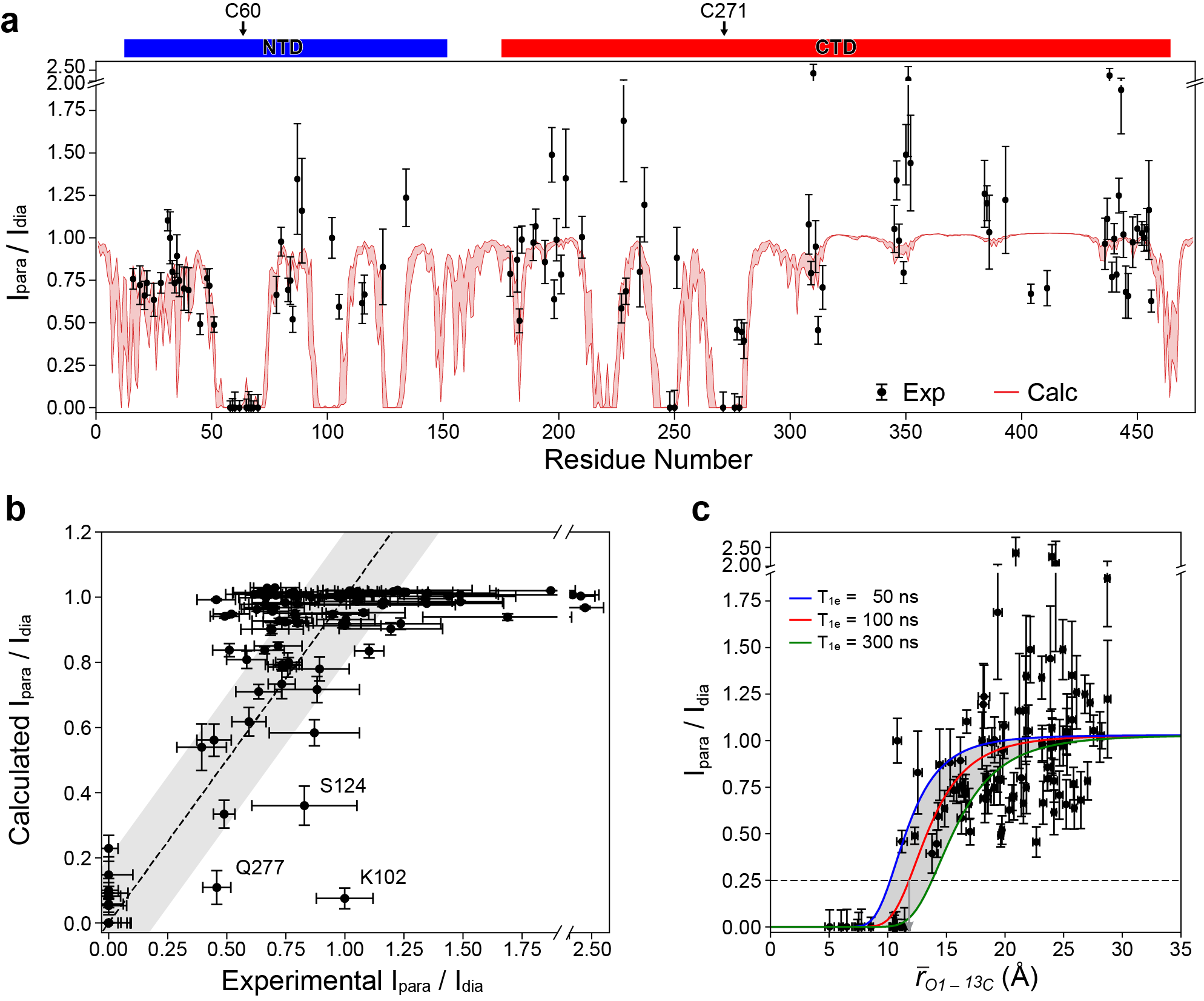

**Figure S5:** *Comparison of experimental and calculated PREs obtained with a 3D NCACB experiment on PROXYL-M-tagged DnaB.* (**a**) Experimentally observed and model-calculated I_para_/I_dia_ signal intensity ratios versus DnaB residue number. The paramagnetic spectrum (I_para_) was measured on PROXYL-M-tagged DnaB and the diamagnetic spectrum (I_dia_) was measured on DnaB with reduced PROXYL-M tag. Vertical bars correspond to estimated errors for experimental and calculated PREs. The latter represent one standard deviation of 100 trials of a Monte Carlo error estimation protocol in which 50% of the model-predicted spin label-protein residue distances were randomly deleted. (**b**) Experimental vs. calculated I_para_/I_dia_ correlation plot. The gray shaded area corresponds to an uncertainty of I_para_/I_dia_ of ±0.2. (**c**) I_para_/I_dia_ ratio versus effective nitroxide-nuclear spin distance for PROXYL-M DnaB. The dashed line indicates the lower limit of the signal intensity ratio below which the signal in the paramagnetic spectrum is usually too weak to be detected. A 25% intensity ratio corresponds to an effective distance of ~12Å, which marks the size of the blind sphere for this tag.

***Discrepancy between observed and calculated PREs for Valine 34 suggests structural flexibility is present in the DnaB NTD.***

We wondered whether the small but noticeable differences between experimentally measured and computationally predicted I_para_/I_dia_ values for the cross peaks of V34, which were unambiguously assigned in 2D spectra, could be due to flexibility in the DnaB structure. In addition to the flexibility of the Gd^3+^-DOTA-M tag (which we modeled by generating conformer ensemble models of this tag) the backbone at and around the tagging site as well as at the loop containing V34 could be flexible, among other regions. Backbone flexibility is not captured by the static X-ray structure model of DnaB, but does exist under the experimental NMR conditions. In order to test this hypothesis, we generated ensembles of backbone conformations for the α1-α2-connecting loop, to which V34 is localized, and for the α2-α3-connecting loop, containing the C60 spin label site, which is spatially close to V34, using the Rosetta backrub protocol. A superimposition of the ensemble of models is shown in Figure S6a. PRE effects induced by Gd^3+^-DOTA-M tags were simulated for the DnaB structural ensemble as described before and the range of I_para_/I_dia_ values for each DnaB residue from this set of simulations was determined (see Figure S6b). The largest I_para_/I_dia_ differences (more than 0.1) across the ensemble of DnaB backrub models were localized to residues in the NTD ring including residues in the α1-α2 (e.g., V34) and α2-α3 loops (colored magenta in Figure S6c). The simulated I_para_/I_dia_ values for V34 varied from 0.47 - 0.6 (Cα-Cβ) or 0.43 - 0.57 (Cα-Cγ_1/2_), respectively, for the ensemble of structures. This demonstrates that V34 and other regions in DnaB are sensitive to backbone flexibility. An even more pronounced effect on I_para_/I_dia_ is expected when the degree of conformational flexibility is increased beyond local backbone motions, e.g., by movements of entire secondary structure elements or domains. An alternative explanation is that the α1-α2 loop was incorrectly modeled in the original X-ray structure model, because the conformation in this region is only poorly defined by the reported low-resolution electron density.

**Rosetta backrub simulation of α1α2 and α2α3 loops of DnaB**

We modeled conformational flexibility in the α1α2 and α2α3 loops using the Rosetta backrub protocol[1, 2], which is computationally much more tractable than running an all-atom molecular dynamics simulation for the large DnaB system. A total of 300 independent Monte Carlo trajectories were performed to generate an ensemble of backbone conformations. Each trajectory used 1500 “backrub” moves to simulate backbone motion. In these moves, short main-chain segments (three to twelve residues) are rotated as rigid bodies about an axis defined by the starting and ending Cα-atom of the segment. K-means clustering was then used to select a set of six distinct DnaB structural models, which differed from each other by a pairwise backbone RMSD of ca. 0.5 Å for the selected loop regions.

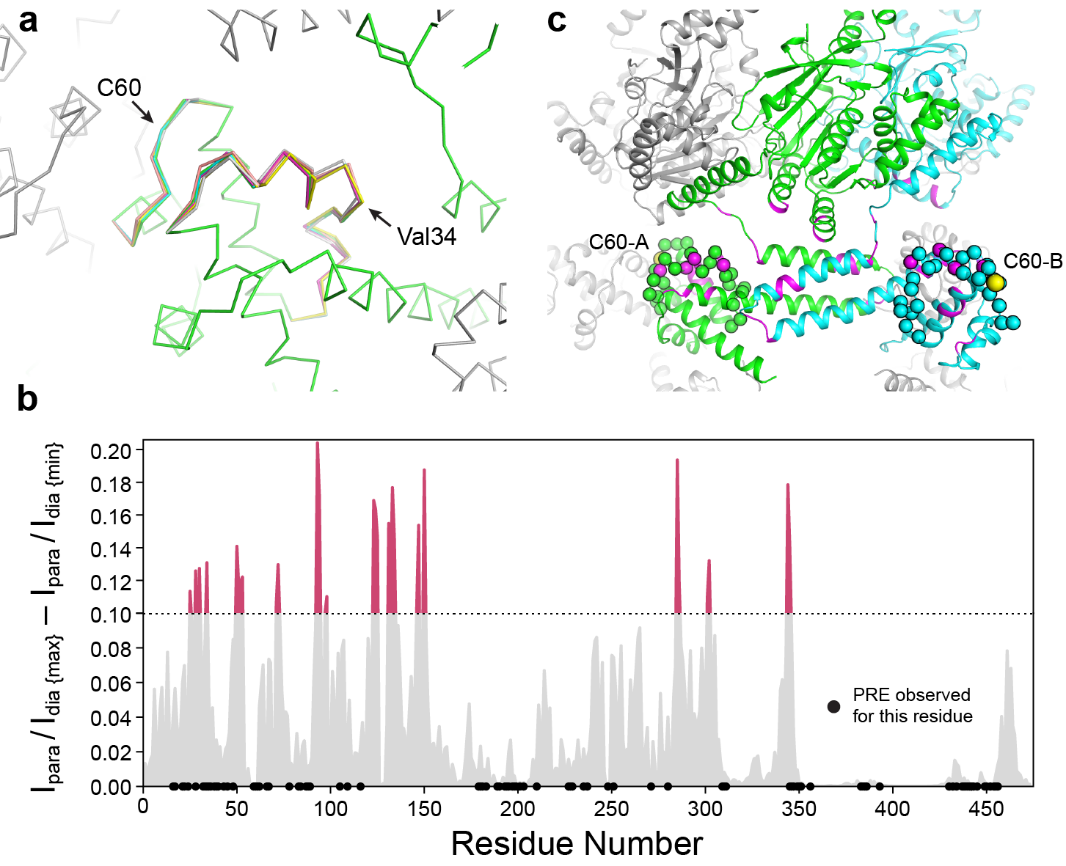

**Figure S6:** *Backrub simulation of α1α2 and α2α3 loops of DnaB, and predicted I_para_/I_dia_ intensity changes for the modeled backrub ensemble.* **(a)** Ensemble of six representative DnaB structure models obtained by a Rosetta backrub simulation of the loop regions connecting helix α1 with α2, and α2 with α3, respectively. The protein Cα-atom trace is shown as ribbon. For each model in the ensemble, the first monomer in shown with a different color, while the neighboring monomers are displayed in gray. The positions of V34 in the α1α2 loop and of C60 in the α2α3 loop are indicated by arrows. **(b)** Difference between the maximal and minimal I_para_/I_dia_ values calculated for the backrub-generated DnaB structure ensemble after computationally tagging the protein with Gd^3+^-DOTA-M. Differences larger than 0.1 are colored magenta. The positions of residues for which an experimental I_para_/I_dia_ value could be measured, are indicated by a dot. **(c)** Residues for which the Ipara/Idia value changes by more than 0.1 (colored magenta) due to the simulated backrub motions are localized to loops α1α2 and α2α3, the NTD ring, and the interface between the NTD and CTD ring. Two monomers of the DnaB structure are colored green and cyan, respectively, while all other monomers are colored gray. Loops α1α2 and α2α3 are shown as spheres. The position of the spin labeled residue C60 is shown with a yellow sphere and labeled.

**Simulation of paramagnetic signal decay profiles for the 2D DARR experiment (Figure S7) and 3D NCACB experiment (Figure S8)**

The signal intensity ratio between para- and diamagnetic spectrum for the 2D DARR experiment was calculated according to:

$\frac{I_{para}}{I_{dia}}=I_{CP}(\tau_{CP})\cdot I_{t1}\cdot I_{DARR}(\tau_{DARR})\cdot I_{t2}$	(S1)

I_CP_ and I_DARR_ are the signal intensity changes during the cross polarization (CP) step and DARR mixing period, respectively. I_t1_ and I_t2_ represent the signal intensity reduction during the evolution (t_1_) and detection (t_2_) periods, respectively. The individual terms in equation (S1) are:

$I_{CP}\left( t \right)=\frac{\exp\left[ \lambda_{1}\left( \Gamma_{1\rho,H},\Gamma_{1\rho,C} \right)t \right]+\exp\left[ \lambda_{2}\left( \Gamma_{1\rho,H},\Gamma_{1\rho,C} \right)t \right]}{1-\exp\left( -2R_{IS}t \right)}$	(S2)

where,	$\lambda_{1}=\frac{-\left( 2R_{IS}+\Gamma_{1\rho,H}+\Gamma_{1\rho,C} \right)+\sqrt{4R_{IS}^{2}+{(\Gamma_{1\rho,H}-\Gamma_{1\rho,C})}^{2}}}{2}$	(S3)

$\lambda_{2}=\frac{-\left( 2R_{IS}+\Gamma_{1\rho,H}+\Gamma_{1\rho,C} \right)-\sqrt{4R_{IS}^{2}+{(\Gamma_{1\rho,H}-\Gamma_{1\rho,C})}^{2}}}{2}$	(S4)

$I_{DARR}\left( t \right)=\exp(-\Gamma_{1,c}t)$	(S5)

$I_{t1}=I_{t2}=\frac{1/\left( R_{2,dia}+\Gamma_{2,C} \right)}{1/R_{2,dia}}=\frac{\pi w_{dia}}{\pi w_{dia}+\Gamma_{2,C}}$	(S6)

$\Gamma_{1\rho,H}$ and $\Gamma_{1\rho,C}$ are the paramagnetic relaxation rates of ^1^H and ^13^C in the rotating frame, $\Gamma_{1,C}$ is the paramagnetic longitudinal relaxation rate of ^13^C, and $\Gamma_{2,C}$ is the paramagnetic transversal relaxation rate of ^13^C. The cross polarization contact time (τ_CP_) was 0.5 ms and the DARR mixing time (τ_DARR_) was 20 ms. The rate constant for the magnetization transfer between ^1^H and ^13^C during the CP step (*R*_IS_) for the experiments with Gd-DOTA-M or PROXYL-M was 6340 s^-1^ or 5770 s^-1^, respectively. Such values were determined from the CP build-up curves obtained upon varying the contact time (see Figure S9, for more details see reference [3]). *w*_dia_ is the line width of the NMR signal in the diamagnetic spectrum.

The intensity ratio of an NMR signal recorded in the para- vs. diamagnetic spectrum of the 3D NCACB experiment was calculated according to:

$\frac{I_{para}}{I_{dia}}=I_{CP1}(\tau_{CP1})\cdot I_{t1}\cdot I_{CP2}(\tau_{CP2})\cdot I_{t2}\cdot I_{DREAM}(\tau_{DREAM})\cdot I_{t3}$	(S7)

where I_CP1_ and I_CP2_ are the change in signal intensity during the cross polarization (CP) steps, I_DREAM_ is the signal reduction during the DREAM mixing period, and I_t1_, I_t2_ and I_t3_ represent the signal intensity reduction during the evolution (t_1_ and t_2_) and detection (t_2_) periods, respectively. The derivation of equation (S7) is described in detail in Tamaki et al. [3]. I_CP1_ and I_CP2_ are dependent on the paramagnetic relaxation rates of ^1^H and ^13^C or ^13^C and ^15^N in the rotating frame (T*_1_*_ρ,H_, T*_1_*_ρ,C_ or T*_1_*_ρ,C_, T*_1_*_ρ,N_), I_DREAM_ is a function of the paramagnetic longitudinal relaxation rate of ^13^C (*T*_1,C_), and I_t1_, I_t2_ and I_t3_ are determined by the paramagnetic transversal relaxation rate of ^13^C (*T*_2,C_) or ^15^N (*T*_2,N_). All used experimental parameters are summarized in Table S4.

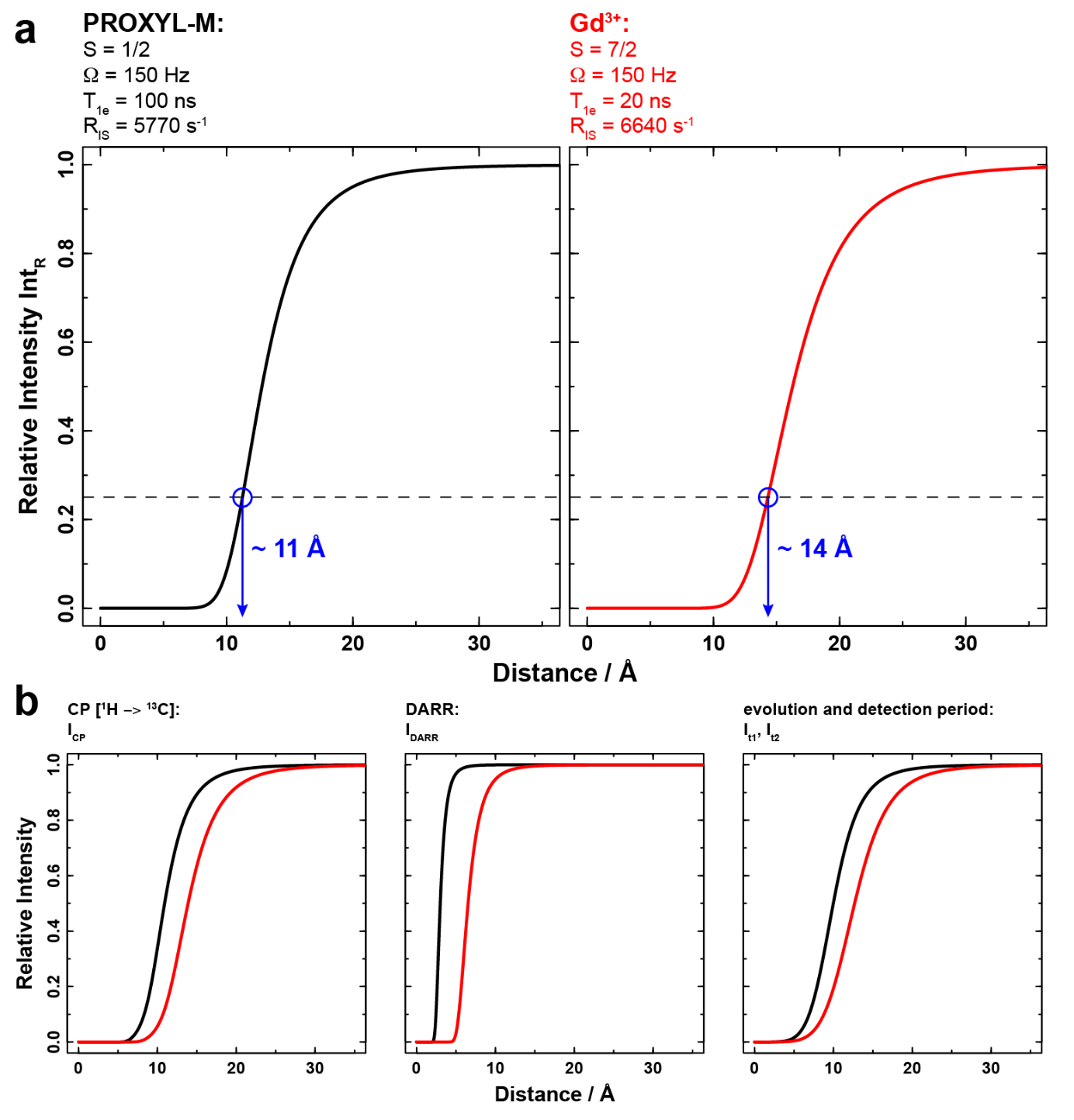

**Figure S7:** *Simulated signal decay profiles for ^13^C-^13^C DARR experiments (with 20ms mixing time) in the presence of PROXYL-M (black) or Gd^3+^-DOTA-M (red) as a function of the electron-nucleus distance.* (**a**) Determination of the size of the blind spheres of nitroxide and Gd^3+^ assuming a lower detection limit of 25 % (dashed line). (**b**) Contributing terms to the total intensity of the signal decay profile: I_CP_, I_DARR_, I_t1_ and I_t2_.

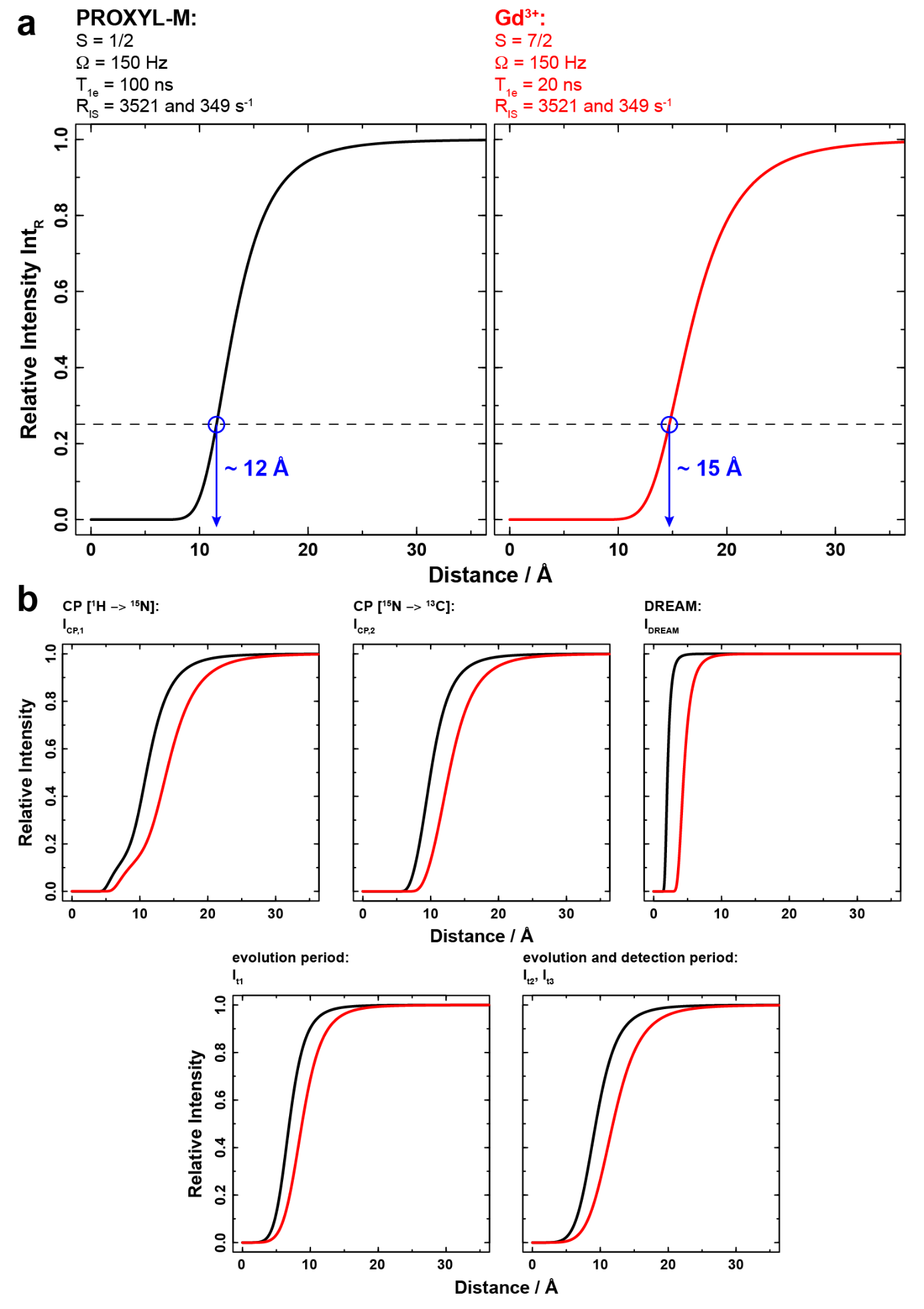

**Figure S8:** *Simulated signal decay profiles of a 3D NCACB experiment in the presence of PROXYL-M (black) or Gd^3+^-DOTA-M (red) as a function of the electron-nucleus distance.* (**a**) Determination of the size of the blind spheres of nitroxide and Gd^3+^ assuming a lower detection limit of 25 % (dashed line). (**b**) Contributing terms to the total intensity of the signal decay profile: I_CP1_, I_CP2_, I_DREAM_, I_t1_, I_t2_ and I_t3_.

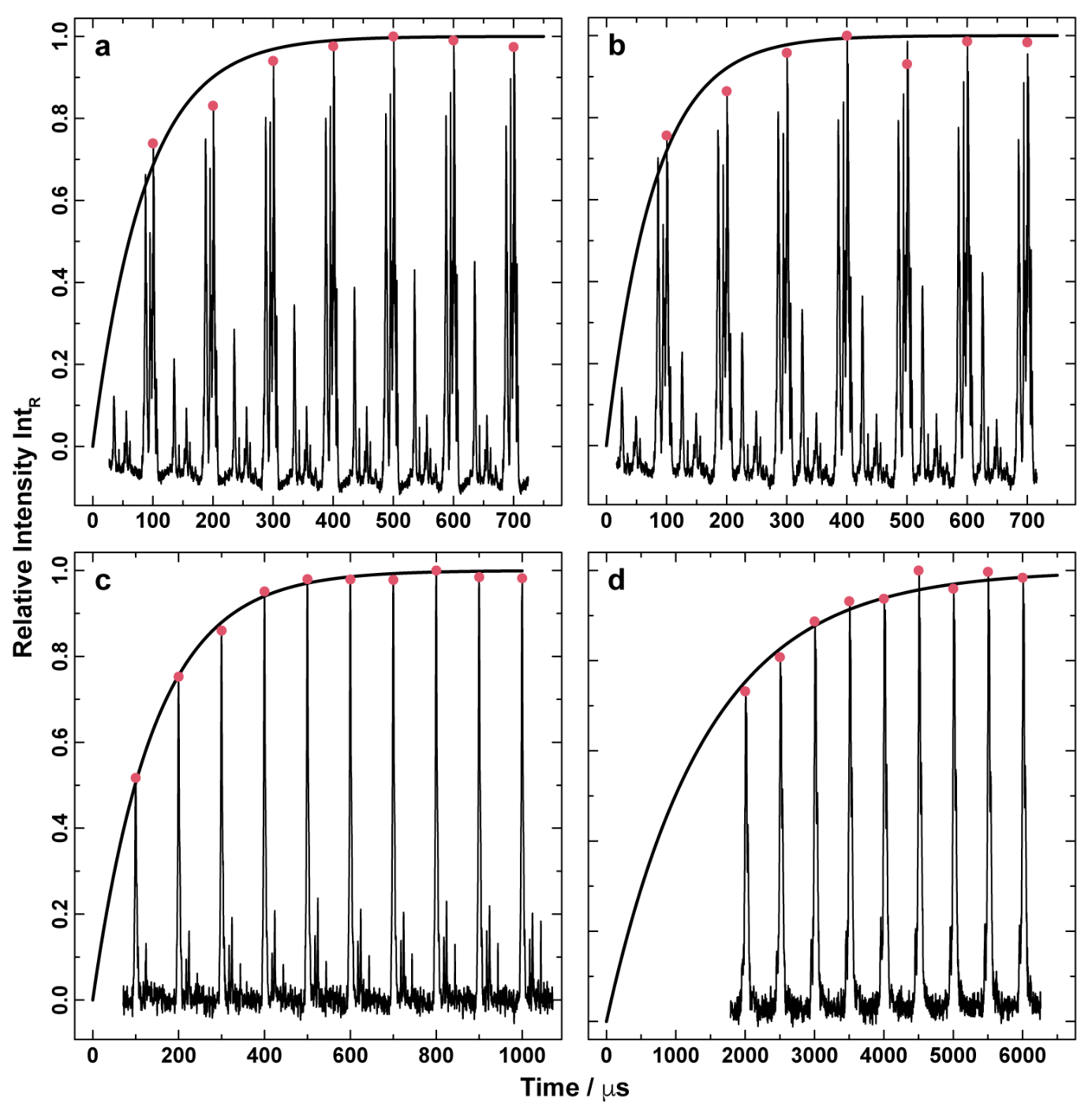

**Figure S9**: *Determination of the rate constant R_IS_ to account for the signal reduction during the CP step.* **a** CP buildup curve for a ^1^H,^13^C CP transfer for the DnaB-PROXYL-M sample, **b** CP buildup curve for a ^1^H,^13^C CP transfer for the DnaB-Gd^3+^-DOTA-M sample, **c** CP buildup curve for a ^1^H,^15^N CP transfer for the DnaB-PROXYL-M sample and **d** CP buildup curve for a ^15^N,^13^C CP transfer for the DnaB-PROXYL-M sample. The straight lines represent the fit to the pink data points (maximum CP signal) according to $I\left( \tau\right)=1-exp(-2R_{IS}\tau)$.

**Table S1:** PRE data (I_para_/I_dia_ ratios) extracted from 2D DARR spectra (with 20 ms mixing time) of DnaB with reduced PROXYL-M tag and PROXYL-M-tagged DnaB. I_para_/I_dia_ values were scaled relative to that one of Asp39. The given error was determined by Gaussian error propagation using the noise level of the spectra as the error for the signal intensities.

| **Amino Acid No.** | **Amino Acid Type** | **Cross Peak** | **I_para_/I_dia_** | **Error** |
| --- | --- | --- | --- | --- |
| 16 | Ile | (Cα, Cβ) | 0.85 | 0.09 |
|  |  | (Cα, Cγ2) | 0.78 | 0.13 |
| 17 | Val | (Cα, Cγb) | 0.92 | 0.14 |
| 21 | Ile | (Cα, Cβ) | 0.95 | 0.11 |
| 22 | Val | (Cα, Cγa) | 0.85 | 0.08 |
|  |  | (Cα, Cγb) | 0.71 | 0.08 |
| 24 | Ala | (Cα, Cβ) | 0.95 | 0.09 |
| 32 | His | (Cα, Cβ) | 0.84 | 0.10 |
| 33 | Ser | (Cα, Cβ) | 1.09 | 0.08 |
| 34 | Val | (Cα, Cβ) | 0.92 | 0.09 |
|  |  | (Cα, Cγa) | 0.96 | 0.12 |
|  |  | (Cα, Cγb) | 0.96 | 0.12 |
| 35 | Leu | (Cα, Cβ) | 0.77 | 0.09 |
| 36 | Glu | (Cα, Cβ) | 0.75 | 0.09 |
| 38 | Ser | (Cα, Cβ) | 1.03 | 0.08 |
| 39 | Asp | (Cα, Cβ) | 1.00 | 0.10 |
| 40 | Phe | (Cα, Cβ) | 1.26 | 0.18 |
| 43 | Pro | (Cα, Cβ) | 0.68 | 0.11 |
| 45 | Asn | (Cα, Cβ) | 0.78 | 0.09 |
| 48 | Phe | (Cα, Cβ) | 0.78 | 0.07 |
| 54 | Lys | (Cα, Cβ) | 0.75 | 0.07 |
| 59 | Asp | (Cα, Cβ) | 0.24 | 0.04 |
| 60 | Cys | (Cα, Cβ) | 0.00 | 0.02 |
| 61 | Pro | (Cα, Cβ) | 0.00 | 0.01 |
| 62 | Ile | (Cα, Cγ2) | 0.00 | 0.04 |
| 66 | Phe | (Cα, Cβ) | 0.00 | 0.05 |
| 67 | Ile | (Cα, Cβ) | 0.00 | 0.05 |
| 78 | Lys | (Cα, Cβ) | 0.81 | 0.08 |
| 83 | Val | (Cα, Cβ) | 0.99 | 0.08 |
|  |  | (Cα, Cγa) | 1.16 | 0.14 |
| 84 | Ala | (Cα, Cβ) | 0.93 | 0.08 |
| 87 | Ala | (Cα, Cβ) | 1.01 | 0.09 |
| 88 | Ala | (Cα, Cβ) | 0.95 | 0.09 |
| 89 | Ser | (Cα, Cβ) | 0.95 | 0.11 |
| 105 | Ser | (Cα, Cβ) | 1.13 | 0.09 |
| 109 | Lys | (Cα, Cβ) | 1.99 | 0.08 |
| 116 | Thr | (Cα, Cβ) | 1.08 | 0.09 |
| 179 | Ile | (Cα, Cβ) | 0.95 | 0.10 |
| 180 | Pro | (Cα, Cβ) | 0.58 | 0.06 |
| 181 | Thr | (Cα, Cβ) | 0.79 | 0.09 |
|  |  | (Cβ, Cγ2) | 0.95 | 0.08 |
|  |  | (Cα, Cγ2) | 0.95 | 0.12 |
| 183 | Phe | (Cα, Cβ) | 0.99 | 0.11 |
| 189 | Tyr | (Cα, Cβ) | 0.81 | 0.09 |
| 190 | Thr | (Cα, Cβ) | 1.08 | 0.11 |
|  |  | (Cβ, Cγ2) | 1.18 | 0.09 |
| 193 | Phe | (Cα, Cβ) | 1.27 | 0.13 |
| 194 | Asn | (Cα, Cβ) | 1.21 | 0.12 |
| 195 | Lys | (Cα, Cβ) | 1.10 | 0.10 |
| 197 | Ser | (Cα, Cβ) | 1.03 | 0.09 |
| 198 | Leu | (Cα, Cβ) | 0.87 | 0.09 |
| 199 | Val | (Cα, Cβ) | 0.86 | 0.09 |
|  |  | (Cα, Cγa) | 1.00 | 0.13 |
| 201 | Ile | (Cα, Cβ) | 1.04 | 0.12 |
| 203 | Ala | (Cα, Cβ) | 1.25 | 0.15 |
| 210 | Thr | (Cα, Cβ) | 1.20 | 0.10 |
|  |  | (Cβ, Cγ2) | 1.12 | 0.09 |
| 227 | Val | (Cα, Cβ) | 0.96 | 0.12 |
|  |  | (Cα, Cγa) | 0.91 | 0.14 |
| 228 | Ala | (Cα, Cβ) | 0.96 | 0.10 |
| 229 | Val | (Cα, Cβ) | 1.02 | 0.13 |
| 235 | Ser | (Cα, Cβ) | 0.90 | 0.09 |
| 237 | Glu | (Cα, Cβ) | 0.79 | 0.12 |
| 248 | Thr | (Cα, Cβ) | 0.00 | 0.03 |
|  |  | (Cβ, Cγ2) | 0.00 | 0.02 |
|  |  | (Cα, Cγ2) | 0.00 | 0.05 |
| 251 | Asn | (Cα, Cβ) | 0.00 | 0.06 |
| 271 | Cys | (Cα, Cβ) | 0.00 | 0.02 |
| 280 | Leu | (Cα, Cβ) | 0.90 | 0.12 |
| 309 | Ile | (Cα, Cγ2) | 1.05 | 0.12 |
| 310 | Ala | (Cα, Cβ) | 1.27 | 0.13 |
| 311 | Phe | (Cα, Cβ) | 1.08 | 0.16 |
| 345 | Leu | (Cα, Cβ) | 0.93 | 0.10 |
| 346 | Glu | (Cα, Cβ) | 1.09 | 0.10 |
| 347 | Ile | (Cα, Cδ1) | NA | NA |
|  |  | (Cα, Cγ2) | 1.10 | 0.13 |
| 349 | Ile | (Cα, Cβ) | 0.94 | 0.09 |
|  |  | (Cα, Cγ2) | 1.09 | 0.15 |
| 350 | Ile | (Cα, Cβ) | 1.14 | 0.14 |
|  |  | (Cα, Cγ2) | 1.26 | 0.20 |
| 351 | Ala | (Cα, Cβ) | 1.45 | 0.15 |
| 354 | Gln | (Cα, Cβ) | 1.17 | 0.21 |
| 356 | Asn | (Cα, Cβ) | 1.10 | 0.12 |
| 383 | Asp | (Cα, Cβ) | 1.06 | 0.10 |
| 384 | Ile | (Cα, Cβ) | 1.13 | 0.13 |
|  |  | (Cα, Cγ2) | 1.00 | 0.10 |
| 385 | Val | (Cα, Cβ) | 1.20 | 0.14 |
|  |  | (Cα, Cγa) | 1.28 | 0.12 |
| 386 | Leu | (Cα, Cβ) | 0.85 | 0.13 |
| 393 | Ile | (Cα, Cβ) | 1.08 | 0.20 |
| 405 | Lys | (Cα, Cβ) | 0.55 | 0.06 |
| 409 | Glu | (Cα, Cβ) | 0.99 | 0.08 |
| 412 | Ile | (Cα, Cβ) | 1.00 | 0.09 |
|  |  | (Cα, Cγ2) | 0.98 | 0.14 |
| 430 | Lys | (Cα, Cβ) | 0.90 | 0.07 |
| 432 | Asn | (Cα, Cβ) | 1.07 | 0.10 |
| 434 | Ser | (Cα, Cβ) | 0.95 | 0.09 |
| 437 | Glu | (Cα, Cβ) | 1.07 | 0.09 |
| 438 | Ala | (Cα, Cβ) | 1.04 | 0.09 |
| 439 | Glu | (Cα, Cβ) | 0.74 | 0.07 |
| 440 | Ile | (Cα, Cγ2) | 0.96 | 0.10 |
| 441 | Ile | (Cα, Cβ) | 0.95 | 0.11 |
|  |  | (Cα, Cγ2) | 0.93 | 0.12 |
| 442 | Val | (Cα, Cγa) | 1.57 | 0.25 |
|  |  | (Cα, Cγb) | 0.93 | 0.08 |
| 443 | Ala | (Cα, Cβ) | 1.05 | 0.09 |
| 445 | Asn | (Cα, Cβ) | 0.78 | 0.13 |
| 449 | Ala | (Cα, Cβ) | 1.34 | 0.11 |
| 450 | Thr | (Cα, Cβ) | 1.04 | 0.11 |
|  |  | (Cα, Cγ2) | 1.28 | 0.14 |
|  |  | (Cβ, Cγ2) | 1.16 | 0.09 |
| 452 | Thr | (Cα, Cβ) | 1.10 | 0.11 |
|  |  | (Cα, Cγ2) | 1.14 | 0.10 |
|  |  | (Cβ, Cγ2) | 1.28 | 0.09 |
| 454 | Tyr | (Cα, Cβ) | 1.18 | 0.14 |
| 455 | Thr | (Cα, Cβ) | 1.05 | 0.15 |
| 456 | Arg | (Cα, Cβ) | 0.86 | 0.08 |

**Table S2:** PRE data (I_para_/I_dia_ ratios) extracted from 2D DARR spectra (with 20 ms mixing time) of DOTA-M-tagged DnaB loaded with either Lu^3+^ or Gd^3+^, respectively. The given error was determined by Gaussian error propagation using the noise level of the spectra as the error for the signal intensities.

| **Amino Acid No.** | **Amino Acid Type** | **Cross Peak** | **I_para_/I_dia_** | **Error** |
| --- | --- | --- | --- | --- |
| 16 | Ile | (Cα, Cβ) | 1.08 | 0.08 |
|  |  | (Cα, Cγ2) | 0.89 | 0.16 |
| 17 | Val | (Cα, Cγb) | 1.02 | 0.14 |
| 21 | Ile | (Cα, Cβ) | 0.77 | 0.09 |
| 22 | Val | (Cα, Cγa) | 0.65 | 0.04 |
|  |  | (Cα, Cγb) | 0.70 | 0.07 |
| 24 | Ala | (Cα, Cβ) | 0.44 | 0.05 |
| 28 | Ile | (Cα, Cδ1) | 0.84 | 0.21 |
| 32 | His | (Cα, Cβ) | 0.76 | 0.09 |
| 33 | Ser | (Cα, Cβ) | 0.87 | 0.02 |
| 34 | Val | (Cα, Cβ) | 0.92 | 0.09 |
|  |  | (Cα, Cγa) | 0.82 | 0.11 |
|  |  | (Cα, Cγb) | 0.82 | 0.11 |
| 35 | Leu | (Cα, Cβ) | 1.14 | 0.11 |
| 36 | Glu | (Cα, Cβ) | 0.63 | 0.09 |
| 38 | Ser | (Cα, Cβ) | 0.81 | 0.04 |
| 39 | Asp | (Cα, Cβ) | 1.09 | 0.12 |
| 40 | Phe | (Cα, Cβ) | 0.75 | 0.13 |
| 43 | Pro | (Cα, Cβ) | 0.72 | 0.13 |
| 45 | Asn | (Cα, Cβ) | 0.73 | 0.10 |
| 48 | Phe | (Cα, Cβ) | 0.78 | 0.05 |
| 54 | Lys | (Cα, Cβ) | 0.69 | 0.02 |
| 59 | Asp | (Cα, Cβ) | 0.00 | NA |
| 60 | Cys | (Cα, Cβ) | 0.00 | NA |
| 61 | Pro | (Cα, Cβ) | 0.00 | NA |
| 62 | Ile | (Cα, Cγ2) | 0.00 | NA |
| 66 | Phe | (Cα, Cβ) | 0.00 | NA |
| 67 | Ile | (Cα, Cβ) | 0.00 | NA |
| 78 | Lys | (Cα, Cβ) | 0.68 | 0.05 |
| 83 | Val | (Cα, Cβ) | 0.86 | 0.04 |
|  |  | (Cα, Cγb) | 0.68 | 0.11 |
| 84 | Ala | (Cα, Cβ) | 0.79 | 0.02 |
| 87 | Ala | (Cα, Cβ) | 0.70 | 0.04 |
| 88 | Ala | (Cα, Cβ) | 0.71 | 0.06 |
| 89 | Ser | (Cα, Cβ) | 0.66 | 0.08 |
| 105 | Ser | (Cα, Cβ) | 1.03 | 0.04 |
| 109 | Lys | (Cα, Cβ) | 0.60 | 0.01 |
| 116 | Thr | (Cα, Cβ) | 0.82 | 0.05 |
| 179 | Ile | (Cα, Cβ) | 0.63 | 0.06 |
| 180 | Pro | (Cα, Cβ) | 0.38 | 0.03 |
| 181 | Thr | (Cα, Cβ) | 0.89 | 0.10 |
|  |  | (Cβ, Cγ2) | 0.57 | 0.03 |
| 183 | Phe | (Cα, Cβ) | 0.47 | 0.08 |
| 189 | Tyr | (Cα, Cβ) | 0.58 | 0.07 |
| 190 | Thr | (Cα, Cβ) | 0.90 | 0.08 |
|  |  | (Cβ, Cγ2) | 0.79 | 0.02 |
| 193 | Phe | (Cα, Cβ) | 0.63 | 0.06 |
| 194 | Asn | (Cα, Cβ) | 0.79 | 0.07 |
| 195 | Lys | (Cα, Cβ) | 0.81 | 0.06 |
| 197 | Ser | (Cα, Cβ) | 0.91 | 0.05 |
| 198 | Leu | (Cα, Cβ) | 1.28 | 0.12 |
| 199 | Val | (Cα, Cβ) | 0.77 | 0.06 |
|  |  | (Cα, Cγa) | 1.04 | 0.16 |
| 201 | Ile | (Cα, Cβ) | 0.85 | 0.10 |
| 203 | Ala | (Cα, Cβ) | 0.92 | 0.09 |
| 210 | Thr | (Cα, Cβ) | 0.82 | 0.05 |
|  |  | (Cβ, Cγ2) | 0.77 | 0.03 |
| 227 | Val | (Cα, Cβ) | 0.65 | 0.09 |
|  |  | (Cα, Cγa) | 0.43 | 0.09 |
| 228 | Ala | (Cα, Cβ) | 0.98 | 0.08 |
| 229 | Val | (Cα, Cβ) | 0.82 | 0.11 |
| 235 | Ser | (Cα, Cβ) | 0.98 | 0.09 |
| 237 | Glu | (Cα, Cβ) | 0.87 | 0.16 |
| 248 | Thr | (Cα, Cβ) | 0.00 | NA |
| 251 | Asn | (Cα, Cβ) | 0.00 | NA |
| 271 | Cys | (Cα, Cβ) | 0.00 | NA |
| 280 | Leu | (Cα, Cβ) | 0.59 | 0.10 |
| 309 | Ile | (Cα, Cδ1) | NA | NA |
|  |  | (Cα, Cγ2) | 1.07 | 0.11 |
| 310 | Ala | (Cα, Cβ) | 0.86 | 0.08 |
| 311 | Phe | (Cα, Cβ) | 1.13 | 0.24 |
| 345 | Leu | (Cα, Cβ) | 0.95 | 0.08 |
| 346 | Glu | (Cα, Cβ) | 0.94 | 0.07 |
| 347 | Ile | (Cα, Cδ1) | NA | NA |
|  |  | (Cα, Cγ1) | NA | NA |
|  |  | (Cα, Cγ2) | 1.96 | 0.24 |
| 349 | Ile | (Cα, Cβ) | 0.92 | 0.07 |
|  |  | (Cα, Cγ2) | 0.90 | 0.12 |
| 350 | Ile | (Cα, Cβ) | 0.90 | 0.12 |
|  |  | (Cα, Cγ2) | 0.95 | 0.18 |
| 351 | Ala | (Cα, Cβ) | 0.96 | 0.08 |
| 356 | Asn | (Cα, Cβ) | 0.82 | 0.10 |
| 383 | Asp | (Cα, Cβ) | 1.13 | 0.10 |
| 384 | Ile | (Cα, Cβ) | 1.03 | 0.11 |
|  |  | (Cα, Cδ1) | 1.46 | 0.38 |
|  |  | (Cα, Cγ2) | 0.97 | 0.07 |
| 385 | Val | (Cα, Cβ) | 1.17 | 0.12 |
|  |  | (Cα, Cγa) | 1.10 | 0.08 |
| 386 | Leu | (Cα, Cβ) | 1.19 | 0.18 |
| 393 | Ile | (Cα, Cβ) | 1.16 | 0.25 |
| 405 | Lys | (Cα, Cβ) | 0.49 | 0.02 |
| 409 | Glu | (Cα, Cβ) | 0.60 | 0.04 |
| 412 | Ile | (Cα, Cβ) | 0.63 | 0.05 |
|  |  | (Cα, Cγ2) | 0.76 | 0.12 |
| 430 | Lys | (Cα, Cβ) | 0.71 | 0.02 |
| 432 | Asn | (Cα, Cβ) | 0.70 | 0.05 |
| 434 | Ser | (Cα, Cβ) | 0.94 | 0.07 |
| 437 | Glu | (Cα, Cβ) | 0.67 | 0.04 |
| 438 | Ala | (Cα, Cβ) | 0.87 | 0.05 |
| 439 | Glu | (Cα, Cβ) | 0.51 | 0.04 |
| 440 | Ile | (Cα, Cγ2) | 0.91 | 0.08 |
| 441 | Ile | (Cα, Cβ) | 0.71 | 0.09 |
|  |  | (Cα, Cγ2) | 0.80 | 0.12 |
| 442 | Val | (Cα, Cγa) | 0.90 | 0.16 |
|  |  | (Cα, Cγb) | 0.77 | 0.04 |
| 443 | Ala | (Cα, Cβ) | 0.88 | 0.05 |
| 445 | Asn | (Cα, Cβ) | 0.46 | 0.12 |
| 449 | Ala | (Cα, Cβ) | 0.73 | 0.05 |
| 450 | Thr | (Cα, Cβ) | 0.75 | 0.08 |
|  |  | (Cα, Cγ2) | 0.80 | 0.09 |
|  |  | (Cβ, Cγ2) | 0.76 | 0.03 |
| 452 | Thr | (Cα, Cβ) | 0.91 | 0.08 |
|  |  | (Cα, Cγ2) | 0.85 | 0.05 |
|  |  | (Cβ, Cγ2) | 0.79 | 0.03 |
| 454 | Tyr | (Cα, Cβ) | 1.05 | 0.12 |
| 455 | Thr | (Cα, Cβ) | 0.75 | 0.14 |
| 456 | Arg | (Cα, Cβ) | 0.66 | 0.04 |

**Table S3:** PRE data (I_para_/I_dia_ ratios) extracted from 3D NCACB spectra of DnaB with reduced PROXYL-M tag and PROXYL-M-tagged DnaB. I_para_/I_dia_ values were scaled relative to that one of His32. The given error was determined by Gaussian error propagation using the noise level of the spectra as the error for the signal intensities.

| **Amino Acid No.** | **Amino Acid Type** | **Cross Peak** | **I_para_/I_dia_** | **Error** |
| --- | --- | --- | --- | --- |
| 16 | Ile | (N, Cα, Cβ) | 0.76 | 0.13 |
| 19 | Ser | (N, Cα, Cβ) | 0.72 | 0.15 |
| 21 | Ile | (N, Cα, Cβ) | 0.66 | 0.13 |
| 22 | Val | (N, Cα, Cβ) | 0.73 | 0.13 |
| 25 | Asn | (N, Cα, Cβ) | 0.64 | 0.14 |
| 28 | Ile | (N, Cα, Cβ) | 0.73 | 0.13 |
| 31 | Val | (N, Cα, Cβ) | 1.10 | 0.15 |
| 32 | His | (N, Cα, Cβ) | 1.00 | 0.20 |
| 33 | Ser | (N, Cα, Cβ) | 0.80 | 0.13 |
| 34 | Val | (N, Cα, Cβ) | 0.73 | 0.13 |
| 35 | Leu | (N, Cα, Cβ) | 0.89 | 0.17 |
| 36 | Glu | (N, Cα, Cβ) | 0.75 | 0.14 |
| 38 | Ser | (N, Cα, Cβ) | 0.70 | 0.16 |
| 40 | Phe | (N, Cα, Cβ) | 0.69 | 0.16 |
| 45 | Asn | (N, Cα, Cβ) | 0.49 | 0.11 |
| 48 | Phe | (N, Cα, Cβ) | 0.76 | 0.13 |
| 49 | Phe | (N, Cα, Cβ) | 0.72 | 0.14 |
| 51 | Ile | (N, Cα, Cβ) | 0.49 | 0.10 |
| 54 | Lys | (N, Cα, Cβ) | 0.00 | 0.02 |
| 58 | Glu | (N, Cα, Cβ) | 0.00 | 0.02 |
| 59 | Asp | (N, Cα, Cβ) | 0.00 | 0.03 |
| 60 | Cys | (N, Cα, Cβ) | 0.00 | 0.05 |
| 62 | Ile | (N, Cα, Cβ) | 0.00 | 0.02 |
| 65 | Asn | (N, Cα, Cβ) | 0.00 | 0.03 |
| 66 | Phe | (N, Cα, Cβ) | 0.00 | 0.06 |
| 67 | Ile | (N, Cα, Cβ) | 0.00 | 0.04 |
| 68 | Arg | (N, Cα, Cβ) | 0.00 | 0.02 |
| 70 | Lys | (N, Cα, Cβ) | 0.00 | 0.05 |
| 78 | Lys | (N, Cα, Cβ) | 0.66 | 0.15 |
| 80 | Glu | (N, Cα, Cβ) | 0.98 | 0.15 |
| 83 | Val | (N, Cα, Cβ) | 0.69 | 0.13 |
| 84 | Ala | (N, Cα, Cβ) | 0.75 | 0.17 |
| 85 | Ile | (N, Cα, Cβ) | 0.52 | 0.12 |
| 87 | Ala | (N, Cα, Cβ) | 1.35 | 0.35 |
| 89 | Ser | (N, Cα, Cβ) | 1.16 | 0.33 |
| 102 | Lys | (N, Cα, Cβ) | 1.00 | 0.17 |
| 105 | Ser | (N, Cα, Cβ) | 0.59 | 0.12 |
| 115 | Asn | (N, Cα, Cβ) | 0.62 | 0.15 |
| 116 | Thr | (N, Cα, Cβ) | 0.67 | 0.15 |
| 124 | Ser | (N, Cα, Cβ) | 0.83 | 0.24 |
| 134 | Ala | (N, Cα, Cβ) | 1.24 | 0.22 |
| 179 | Ile | (N, Cα, Cβ) | 0.79 | 0.17 |
| 182 | Gly | (N, Cα, Cα) | 0.87 | 0.22 |
| 183 | Phe | (N, Cα, Cβ) | 0.51 | 0.11 |
| 184 | Val | (N, Cα, Cβ) | 0.99 | 0.15 |
| 189 | Tyr | (N, Cα, Cβ) | 0.97 | 0.16 |
| 190 | Thr | (N, Cα, Cβ) | 1.07 | 0.17 |
| 194 | Asn | (N, Cα, Cα) | 0.86 | 0.17 |
| 197 | Ser | (N, Cα, Cβ) | 1.49 | 0.23 |
| 198 | Leu | (N, Cα, Cβ) | 0.64 | 0.15 |
| 199 | Val | (N, Cα, Cβ) | 0.99 | 0.18 |
| 201 | Ile | (N, Cα, Cβ) | 0.78 | 0.16 |
| 203 | Ala | (N, Cα, Cβ) | 1.35 | 0.32 |
| 210 | Thr | (N, Cα, Cβ) | 1.00 | 0.18 |
| 227 | Val | (N, Cα, Cβ) | 0.58 | 0.13 |
| 228 | Ala | (N, Cα, Cβ) | 1.69 | 0.39 |
| 229 | Val | (N, Cα, Cβ) | 0.69 | 0.13 |
| 235 | Ser | (N, Cα, Cβ) | 0.80 | 0.22 |
| 237 | Glu | (N, Cα, Cβ) | 1.19 | 0.25 |
| 248 | Thr | (N, Cα, Cβ) | 0.00 | 0.05 |
| 250 | Ile | (N, Cα, Cβ) | 0.00 | 0.06 |
| 251 | Asn | (N, Cα, Cβ) | 0.88 | 0.21 |
| 271 | Cys | (N, Cα, Cβ) | 0.00 | 0.04 |
| 276 | Ser | (N, Cα, Cβ) | 0.00 | 0.05 |
| 277 | Gln | (N, Cα, Cβ) | 0.46 | 0.10 |
| 278 | Lys | (N, Cα, Cβ) | 0.00 | 0.04 |
| 279 | Lys | (N, Cα, Cβ) | 0.45 | 0.11 |
| 280 | Leu | (N, Cα, Cβ) | 0.39 | 0.12 |
| 308 | Gly | (N, Cα, Cα) | 1.08 | 0.22 |
| 309 | Ile | (N, Cα, Cβ) | 0.79 | 0.13 |
| 310 | Ala | (N, Cα, Cβ) | 2.35 | 0.46 |
| 311 | Phe | (N, Cα, Cβ) | 0.95 | 0.19 |
| 312 | Ile | (N, Cα, Cβ) | 0.46 | 0.11 |
| 314 | Tyr | (N, Cα, Cβ) | 0.71 | 0.16 |
| 345 | Leu | (N, Cα, Cβ) | 1.05 | 0.19 |
| 346 | Glu | (N, Cα, Cβ) | 1.34 | 0.19 |
| 347 | Ile | (N, Cα, Cβ) | 0.98 | 0.16 |
| 349 | Ile | (N, Cα, Cβ) | 0.80 | 0.13 |
| 350 | Ile | (N, Cα, Cβ) | 1.49 | 0.24 |
| 351 | Ala | (N, Cα, Cβ) | 2.07 | 0.61 |
| 352 | Leu | (N, Cα, Cβ) | 1.44 | 0.32 |
| 384 | Ile | (N, Cα, Cβ) | 1.26 | 0.24 |
| 385 | Val | (N, Cα, Cβ) | 1.20 | 0.18 |
| 386 | Leu | (N, Cα, Cβ) | 1.03 | 0.25 |
| 393 | Ile | (N, Cα, Cβ) | 1.22 | 0.34 |
| 404 | Asp | (N, Cα, Cβ) | 0.67 | 0.12 |
| 409 | Glu | (N, Cα, Cβ) | 0.84 | 0.14 |
| 411 | Lys | (N, Cα, Cβ) | 0.70 | 0.15 |
| 412 | Ile | (N, Cα, Cβ) | 0.79 | 0.13 |
| 436 | Glu | (N, Cα, Cβ) | 0.97 | 0.19 |
| 437 | Glu | (N, Cα, Cβ) | 1.11 | 0.18 |
| 438 | Ala | (N, Cα, Cβ) | 2.25 | 0.37 |
| 439 | Glu | (N, Cα, Cβ) | 0.77 | 0.14 |
| 440 | Ile | (N, Cα, Cβ) | 0.99 | 0.16 |
| 441 | Ile | (N, Cα, Cβ) | 0.78 | 0.15 |
| 442 | Val | (N, Cα, Cβ) | 1.25 | 0.18 |
| 443 | Ala | (N, Cα, Cβ) | 1.87 | 0.31 |
| 444 | Lys | (N, Cα, Cβ) | 1.02 | 0.19 |
| 445 | Asn | (N, Cα, Cβ) | 0.68 | 0.17 |
| 446 | Arg | (N, Cα, Cβ) | 0.66 | 0.16 |
| 448 | Gly | (N, Cα, Cα) | 0.97 | 0.19 |
| 450 | Thr | (N, Cα, Cβ) | 1.05 | 0.16 |
| 452 | Thr | (N, Cα, Cβ) | 1.03 | 0.15 |
| 453 | Val | (N, Cα, Cβ) | 1.00 | 0.15 |
| 454 | Tyr | (N, Cα, Cβ) | 1.05 | 0.18 |
| 455 | Thr | (N, Cα, Cβ) | 1.16 | 0.31 |
| 456 | Arg | (N, Cα, Cβ) | 0.63 | 0.12 |

**Table S4**: Overview of the parameters used for the solid-state NMR experiments. For more details about the used adiabatic CP steps and the tangential shapes see reference [4].

| **Experiment** | **2D ^13^C-^13^C 20 ms DARR *Hp*DnaB PROXYL-M red.** | **2D ^13^C-^13^C 20 ms DARR *Hp*DnaB PROXYL-M** | **2D ^13^C-^13^C 20 ms DARR *Hp*DnaB DOTA Lu^3+^** |
| --- | --- | --- | --- |
| ν_r_ / kHz | 17 | 17 | 17 |
| *B*_0_/ T | 20 | 20 | 20 |
| transfer I | HC-CP | HC-CP | HC-CP |
| *ν*_1_(^1^H)/ kHz | 60 | 60 | 60 |
| *ν*_1_(^13^C)/ kHz | 42.6 | 40.4 | 42.7 |
| Shape | Tangent ^1^H | Tangent ^1^H | Tangent ^1^H |
| ^13^C carrier / ppm | 100 | 100 | 100 |
| time / ms | 0.6 | 0.5 | 0.5 |
| transfer II | DARR | DARR | DARR |
| *ν*_1_(^1^H)/ kHz | 17 | 17 | 17 |
| ^13^C carrier / ppm | 100 | 100 | 100 |
| time / ms | 20 | 20 | 20 |
| *t_1_* increments | 2560 | 2560 | 2560 |
| sweep width (*t_1_*) / kHz | 100 | 100 | 100 |
| acquisition time (*t_1_*) / ms | 12.8 | 12.8 | 12.8 |
| *t_2_* increments | 3072 | 3072 | 3072 |
| sweep width (*t_2_*) / kHz | 100 | 100 | 100 |
| acquisition time (*t_2_*) / ms | 15.4 | 15.4 | 15.4 |
| FID resolution F1 / ppm | 0.37 | 0.37 | 0.37 |
| FID resolution F2 / ppm | 0.30 | 0.30 | 0.30 |
| ^1^H Spinal64 decoupling power / kHz | 90 | 90 | 90 |
| interscan delay / s | 2.7 | 2.5 | 2.5 |
| number of scans | 24 | 24 | 24 |
| measurement time / h | 48 | 44 | 44 |

**Table S4 continued.**

| **Experiment** | **2D ^13^C-^13^C 20 ms DARR *Hp*DnaB DOTA Gd^3+^** | **3D NCACB *Hp*DnaB PROXYL-M red.** | **3D NCACB *Hp*DnaB PROXYL-M** |
| --- | --- | --- | --- |
| ν_r_ / kHz | 17 | 17 | 17 |
| *B*_0_/ T | 20 | 20 | 20 |
| transfer I | HC-CP | HN-CP | HN-CP |
| *ν*_1_(^1^H)/ kHz | 60 | 60 | 60 |
| *ν*_1_(^13^C)/ kHz | 40 | - | - |
| *ν*_1_(^15^N)/ kHz | - | 41.9 | 42.9 |
| Shape | Tangent ^1^H | Tangent ^1^H | Tangent ^1^H |
| ^13^C carrier / ppm | 100 | - | - |
| ^15^N carrier / ppm | - | 120 | 120 |
| time / ms | 0.5 | 0.65 | 0.6 |
| transfer II | DARR | NC-CP | NC-CP |
| *ν*_1_(^1^H)/ kHz | 17 | - | - |
| *ν*_1_(^13^C)/ kHz | - | 6 | 6 |
| *ν*_1_(^15^N)/ kHz | - | 10.6 | 10.4 |
| Shape | - | Tangent ^13^C | Tangent ^13^C |
| ^13^C carrier / ppm | 100 | 57 | 57 |
| time / ms | 20 | 5.0 | 4.5 |
| transfer III | - | DREAM | DREAM |
| *ν*_1_(^13^C)/ kHz | - | 8.4 | 7.2 |
| Shape | - | Tangent ^13^C | Tangent ^13^C |
| ^13^C carrier / ppm | - | 53 | 53 |
| time / ms | - | 1.5 | 2 |
| *t_1_* increments | 2560 | 70 | 62 |
| sweep width (*t_1_*) / kHz | 100 | 6 | 6 |
| acquisition time (*t_1_*) / ms | 12.8 | 5.8 | 5.1 |
| *t_2_* increments | 3072 | 132 | 128 |
| sweep width (*t_2_*) / kHz | 100 | 8.6 | 8.6 |
| acquisition time (*t_2_*) / ms | 15.4 | 7.7 | 7.5 |
| *t_3_* increments | - | 2304 | 2304 |
| sweep width (*t_3_*) / kHz | - | 100 | 100 |
| acquisition time (*t_3_*) / ms | - | 11.5 | 11.5 |
| FID resolution F1 / ppm | 0.37 | 2.00 | 2.26 |
| FID resolution F2 / ppm | 0.30 | 0.61 | 0.63 |
| FID resolution F3 / ppm | - | 0.41 | 0.41 |
| ^1^H Spinal64 decoupling power / kHz | 90 | 90 | 90 |
| interscan delay / s | 2.5 | 2.7 | 2.7 |
| number of scans | 40 | 16 | 24 |
| measurement time / h | 73 | 113 | 144 |

**References**

[1]	G.D. Friedland, A.J. Linares, C.A. Smith, T. Kortemme, A Simple Model of Backbone Flexibility Improves Modeling of Side-chain Conformational Variability, J. Mol. Biol., 380 (2008) 757-774.

[2]	I.W. Davis, W.B. Arendall, D.C. Richardson, J.S. Richardson, The Backrub Motion: How Protein Backbone Shrugs When a Sidechain Dances, Structure, 14 (2006) 265-274.

[3]	H. Tamaki, A. Egawa, K. Kido, T. Kameda, M. Kamiya, T. Kikukawa, T. Aizawa, T. Fujiwara, M. Demura, Structure determination of uniformly 13C, 15N labeled protein using qualitative distance restraints from MAS solid-state 13C-NMR observed paramagnetic relaxation enhancement, J. Biomol. NMR, 64 (2016) 87-101.

[4]	S. Hediger, B.H. Meier, N.D. Kurur, G. Bodenhausen, R.R. Ernst, Nmr Cross-Polarization by Adiabatic Passage through the Hartmann-Hahn Condition (Aphh), Chem Phys Lett, 223 (1994) 283-288.
